# Phylogenetic variation of immature neurons in mammalian amygdala: high prevalence in primate expanded nuclei projecting to neocortex

**DOI:** 10.1101/2024.12.27.630481

**Authors:** Marco Ghibaudi, Chiara La Rosa, Nikita Telitsyn, Jean-Marie Graїc, Chris G. Faulkes, Chet C. Sherwood, Luca Bonfanti

**Affiliations:** Neuroscience Institute Cavalieri Ottolenghi (NICO), Orbassano, Italy; Department of Veterinary Sciences, University of Turin, Torino, Italy; Department of Comparative Biomedicine and Food Science, University of Padova, Legnaro, Padova, Italy; School of Biological and Behavioural Sciences, Fogg Building, Mile End Road, Queen Mary University of London, London E1 4NS, UK; Department of Anthropology and Center for the Advanced Study of Human Paleobiology, The George Washington University, Washington DC, USA

## Abstract

Structural changes involving new neurons can occur through stem cell-driven neurogenesis and late-maturing “immature” neurons, namely undifferentiated neuronal precursors frozen in a state of arrested maturation. The latter exist in the cerebral cortex, being particularly abundant in large-brained mammals. Similar cells have been described in the amygdala of some species, although their interspecies variation remain poorly understood. Here, their occurrence, number, molecular expression, and morphology were systematically analyzed in eight diverse mammalian species widely differing in neuroanatomy, brain size, lifespan, and socioecology. We show remarkable phylogenetic variation of the immature neurons in the amygdala, with a significantly greater prevalence in primates. The cells are associated with the amygdala’s basolateral complex that in evolution has expanded in primates in conjunction with cortical projections, thus mimicking the general trend of the neocortex. These results support the emerging view that large brains performing complex socio-cognitive functions rely on wide reservoirs of immature neurons.

## Introduction

In recent years, new interest in brain plasticity has been raised by observations of populations of “immature” or “dormant” neurons (INs) consisting of prenatally generated neuronal cells that can remain in a state of arrested maturation for long time (1–7). These cells do not divide in postnatal or adult life, yet they express the same neuronal markers of immaturity generated by stem cell division in the canonical neurogenic sites (1, 8, 9), e.g., the cytoskeletal protein doublecortin (DCX; 10, 11), and the polysialylated form of the neural cell adhesion molecule (PSA-NCAM; 5, 9). These markers can be expressed for a long time before the cells restart maturation to functionally integrate into neural circuits, in a sort of “neurogenesis without division” (12–14). These kinds of neurons have been characterized in most detail in layer II of the piriform cortex (cortical immature neurons; cINs), by following their progressive maturation in a DCX-Cre-ERT2/Flox-EGFP transgenic mouse model (6, 8, 12). Remarkable interspecies differences of the cINs have also been revealed by systematic quantitative analysis carried out in the entire cerebral cortex of widely different mammals (15). A far higher cell density of cINs is present in large-brained, gyrencephalic species with respect to small-brained, lissencephalic ones, thus suggesting that neurons in arrested maturation might have been increased across evolution to support complex cortical cognitive functions in mammals with reduced stem-cell driven neurogenesis (16, 17). Accordingly, thousands of DCX^+^ immature neurons are present in cortical layer II of humans, extending into their widely expanded neocortex (18–20).

Over the years, the existence of neurons expressing DCX and PSA-NCAM has been repeatedly reported in subcortical brain regions, including amygdala, claustrum, and white matter tracts (21–32). Due to their shared expression of immaturity markers with newborn neuroblasts (9, 32), these cell populations were sometimes considered to be a possible product of non-canonical (parenchymal) neurogenesis (see for example 21, 27), though definitive evidence for their active division was elusive (26, 29, 30, 32–34). Several reports on DCX^+^ neurons in amygdala have included non-rodent mammals, suggesting the existence of interspecies variation (24, 26, 29, 30). A recent study revealed a small population of prenatally generated DCX^+^ cells in the mouse amygdala, mostly restricted to a very thin, previously undetected, paralaminar nucleus (7). The same research group also described the occurrence of high amounts of similar cells in the paralaminar nucleus of humans (30). These data suggest that the occurrence of subcortical immature neurons (scINs) like those in the cortex might be a conserved trait, yet with interspecies differences.

At present, the picture is fragmentary, due to the novelty of the field and because of different methods employed by researchers in the study of single species, often leading to divergent interpretations of the results (32, 35). On this basis, we undertook a systematic, interspecies investigation of the amygdala of diverse mammals by using the same approach employed for assessing the phylogenetic variation of cINs (15). Eight mammalian species that widely differ for neuroanatomy (brain size, gyrencephaly, encephalization) and other life history and socioecological features (lifespan, habitat, food habit) were considered (Fig. 1). They include two rodents characterized by short and long lifespans (mouse and naked mole rat, respectively), a small platyrrhine monkey primate (marmoset) and a large hominoid primate (chimpanzee), a very large brain-sized herbivorous perissodactyl (horse), a carnivoran species showing the highest IN density in the cerebral cortex (cat; 15), the rabbit, whose cIN density is intermediate between small-brained and large-brained mammals, and the sheep, as a gyrencephalic artiodactyl species in which INs of the amygdala where shown to be prenatally generated (29). Different age groups were also investigated, reaching a total of 80 brains analyzed (Fig. 1 and Table S1). This study addresses open questions concerning: i) whether these cells in the amygdala share common features of non-dividing “immature” neurons across species; ii) to what extent they display phylogenetic variation; iii) whether their amount varies with age and how this differs among species; and iv) whether they show similarities with their counterpart in the cortex. DCX^+^ (immature) neurons and Ki67^+^ (dividing) cells were studied to describe their occurrence, morphological and phenotypic types, density, and topographical distribution, and analyzed in a phylogenetic perspective.

**Figure 1.**
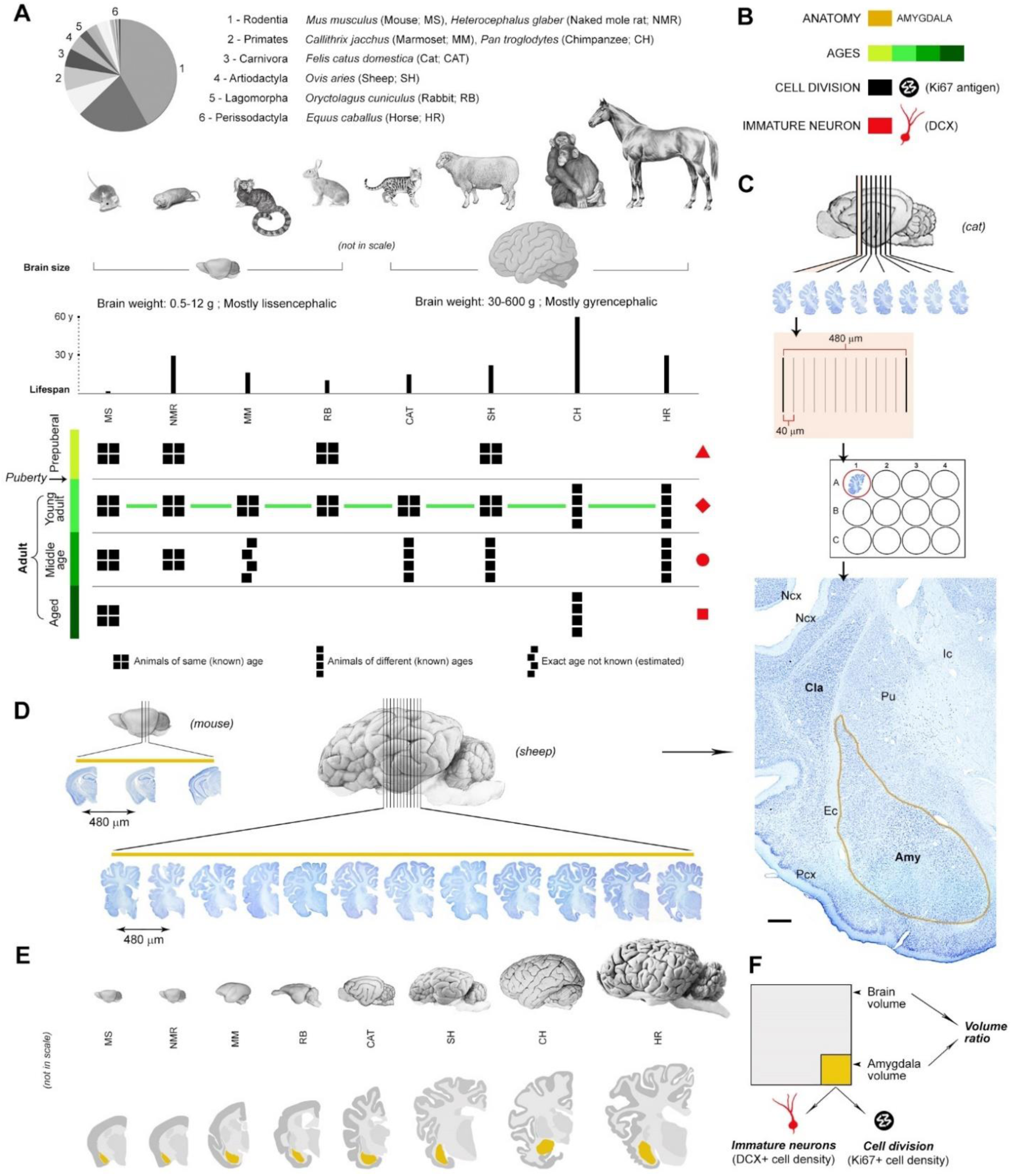
Sample of species, ages, and comparable brain regions of the mammals used in this study. (further information in Tables S1; Animal species are arranged from left to right according to their brain size). (**A**, top) mammalian species and orders (scientific name, common name – used hereafter – and abbreviation) with special reference to their brain size and lifespan. (**A**, bottom) different ages considered for each species, from prepuberal to old aged; all groups are composed of four individuals (black squares), and all species are available at the young adult stage (green line); (Table S1). (**B**) Color code. (**C**,**D**) Brain tissue processing adopted to obtain comparable data in all species considered (mouse, sheep, and cat are represented as an example); serial coronal sections 40 µm thick of the entire hemisphere of each animal species were placed in multi-well plates to have an interval of 480 µm in each well to analyze the anterior-posterior extension of amygdala, followed by staining of sections and segmentation of the subcortical region based on histology (**B**, bottom); the final drawings of neuroanatomy in coronal sections are represented in **E**). (**D**) Different numbers of sections were obtained in each species depending on the brain size and consequent extension of the amygdala (see also Fig. S1 and Table S4). (**F**) By using the comparable method described above, volumes of the amygdala and whole hemisphere were calculated in each species. Then, counting of DCX^+^ cells (immature-appearing neurons) and Ki67^+^ nuclei (dividing cells) were performed in the amygdala (see Fig. 4). Scale bar: 1000 µm.

## Results

### Characterization of the amygdala DCX^+^ neuronal population across mammals

The entire the amygdala of 8 mammalian species was examined with immunodetection of a panel of antigens (Table S2). After testing immunocytochemical staining for the best performing antibodies to detect DCX and Ki67 antigens, the polyclonal goat anti-DCX antibody from Santa Cruz and the polyclonal rabbit anti-Ki67 antibody from Abcam were selected (19; Table S2). Despite slight differences in postmortem interval, fixation procedure, and fixation time (Table S2), no substantial variation was observed in the quality and intensity of the staining (Fig. 2). In addition to the amygdala as a region of interest, other regions from the same brains were used as internal controls, including the canonical neurogenic sites (forebrain subventricular zone, SVZ, and hippocampal subgranular zone, SGZ; Fig. 3B) as a positive control for cell division, and the cerebral cortex (neocortex and paleocortex; Fig. 3B), as a positive control for cINs and negative control for their possible cell division (absence of DCX/Ki67 co-expression).

**Figure 2.**
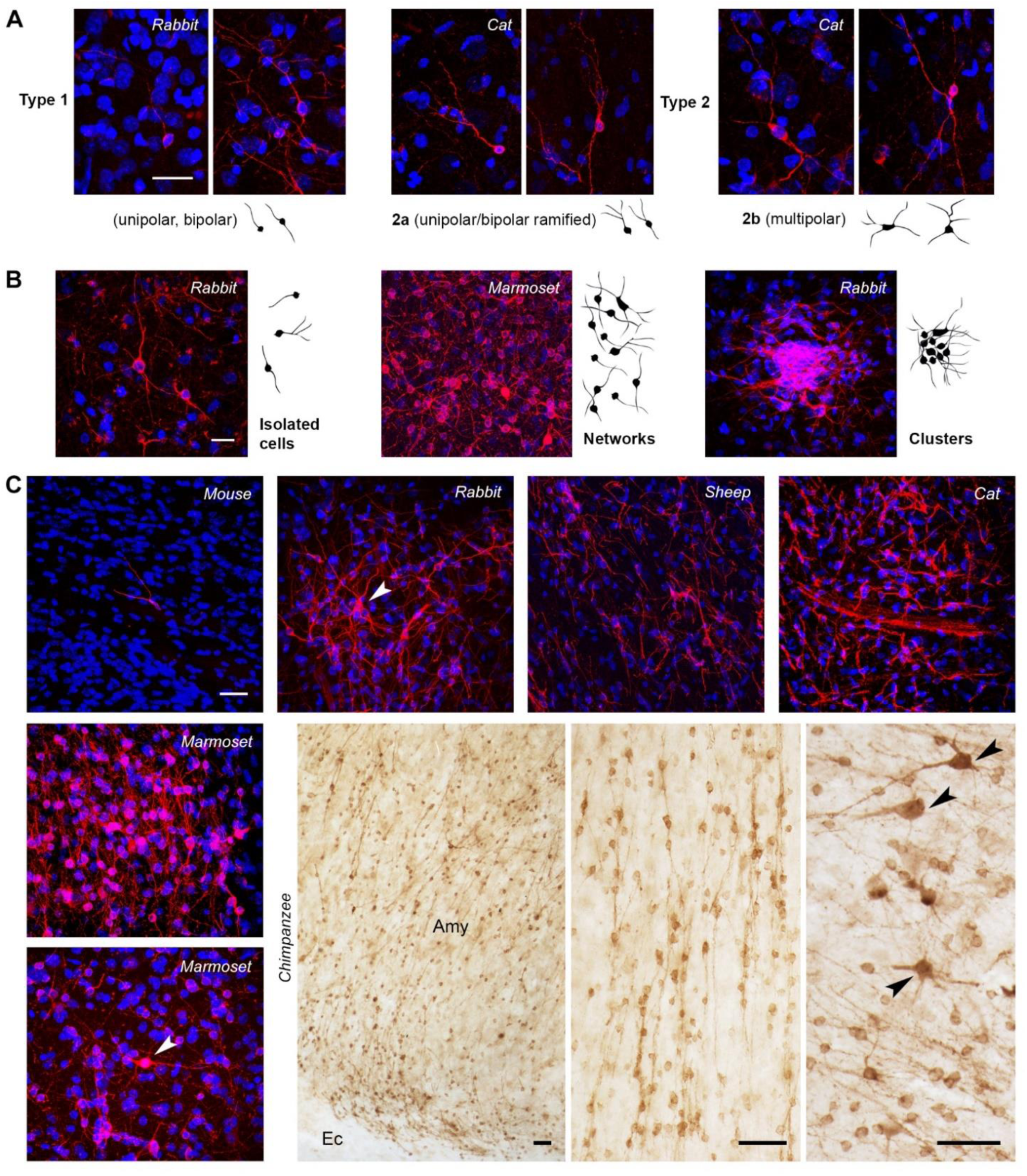
Occurrence, morphology, and general distribution of DCX^+^ cells in the amygdala of mammals. (A) In all species considered, the morphological cell types of DCX^+^ cells were reminiscent of type 1 (very simple morphology) and type 2 (complex cells) immature or dormant neurons described in the cortex (8, 15). (B) At least three types of cell distribution/aggregation were observed, from isolated cells to tightly packed clusters. (**C**) Confocal fields of DCX^+^ cells in the amygdala of different mammals after immunofluorescence staining (except for chimpanzee specimens, photographed in light microscopy after DAB staining). Note the presence of scattered, isolated immunoreactive cells in mouse with respect to dense, extensive networks in primates. Amy, amygdala; Ec, external capsule; arrowheads, examples of type 2b cells. Scale bars: A-C (confocal), 30 µm; C (DAB), 50 µm.

**Figure 3.**
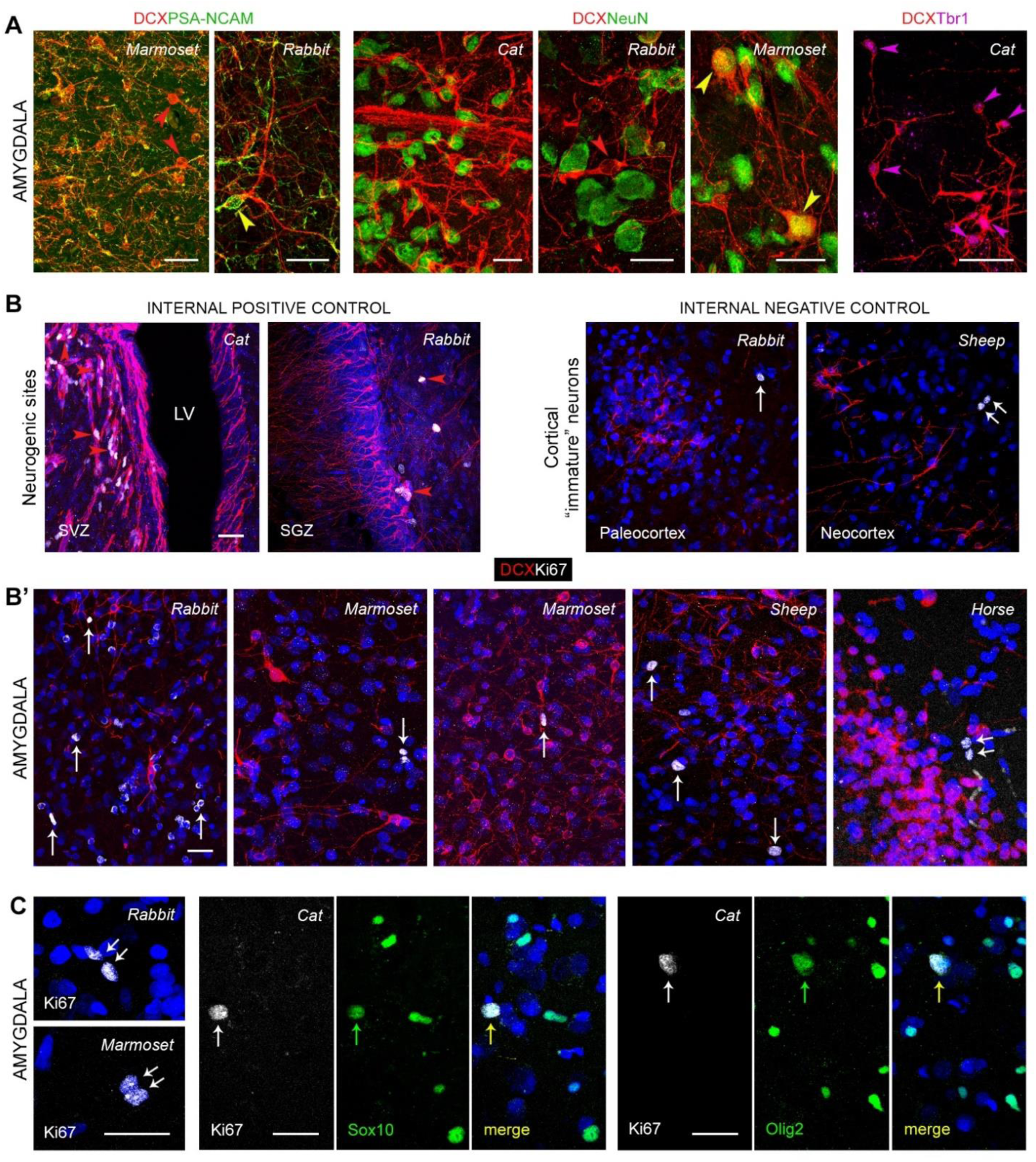
Confocal analysis of a panel of different markers in the amygdala of mammals. (**A**) Markers of immaturity (DCX and PSA-NCAM, left) are widely distributed and co-expressed in the scIN population; some DCX^+^ type 2 cells are devoid of PSA-NCAM (red arrowheads), while others still express it (yellow arrowhead). Most DCX^+^ cells, including type 2 cells (red arrowhead), do not express the marker for postmitotic neurons that start differentiation NeuN (middle); only a subpopulation of type 2 cells express NeuN, indicating they started the maturation process (yellow arrowheads). Most DCX^+^ cells also co-express the marker for glutamatergic neurons Tbr1 (right). (**B**) Positive (neurogenic sites: subventricular zone, SVZ, and subgranular zone, SGZ; LV, lateral ventricle) and negative (cerebral cortex) controls for detection of cell division. (**B’**) Representative images of DCX/Ki67 antigen double staining showing total absence of co-expression in amygdala of any of the species considered. Note the occurrence of “douplets” (double arrow) in the cortical and amygdalar parenchyma. (**C**) Ki67 antigen staining in amygdala frequently revealed “douplets” (left), and was usually associated with oligodendrocyte progenitor cell division; accordingly, frequent co-expression was detectable in double staining of Ki67 antigen with the glial markers SOX10 and Olig2. Scale bars: 30 µm.

DCX^+^ cells were found in the amygdala of all mammals studied, and at all ages in the sample, yet with evident interspecies differences in their density and distribution. Most strikingly, only a few, scattered DCX^+^ cells were detectable in adult mice (being absent in all the coronal sections analyzed), while extended networks of DCX^+^ cells were present in the amygdala of primates (Fig. 2).

Two morphological DCX^+^ cell types previously reported in cortical layer II (cINs of both paleo and neocortex; 15, 29; Figs. 2A and 4B) were consistently detected: type 1, unipolar or bipolar, with a small cell soma (cell soma diameter range: 3-9 μm), and type 2, characterized by ramified dendrites and a larger cell soma (9-19 μm). Type 1 cells were very similar to the correspondent type in the cortex, while type 2 cells were split into two subtypes distinctive to the amygdala: type 2a (bipolar, with a ramified dendrite; soma size range 9-12 µm) and type 2b (showing multiple, ramified dendrites resulting in a multipolar morphology, and 12-19 μm soma size range; Fig. 2A and 4B). In cortical layer II, type 1 and 2 cells have been recognized to represent different stages of maturation in a process leading the small (highly immature) neurons to become larger and more ramified (complex cells) after they restart the maturation process to become principal neurons of that layer (pyramidal neurons; 6, 8, 12). Accordingly, the large, type 2b cells described here resemble the principal cell type of the amygdala (pyramidal-like Class I projection neurons; 36, 37), and their glutamatergic identity was confirmed by co-expression with the excitatory neuron transcription factor T-box brain 1 (Tbr1; 38, 39; Fig. 3A). A small subpopulation of the DCX^+^ cells (percentages given in Fig. 5C’), consisting of large, complex cells, mostly type 2b cells, also co-expressed NeuN (Fig. 3A and 5C; tabulated data can be found in data S1), indicating they have started maturation. On the other hand, most of the DCX^+^ cells, including all type 1 cells and the remaining Type 2 cells, consistently co-expressed the “low adhesive” adhesion molecule PSA-NCAM on their soma and/or cell processes (Fig. 3A), thus confirming their immaturity state.

**Figure 4.**
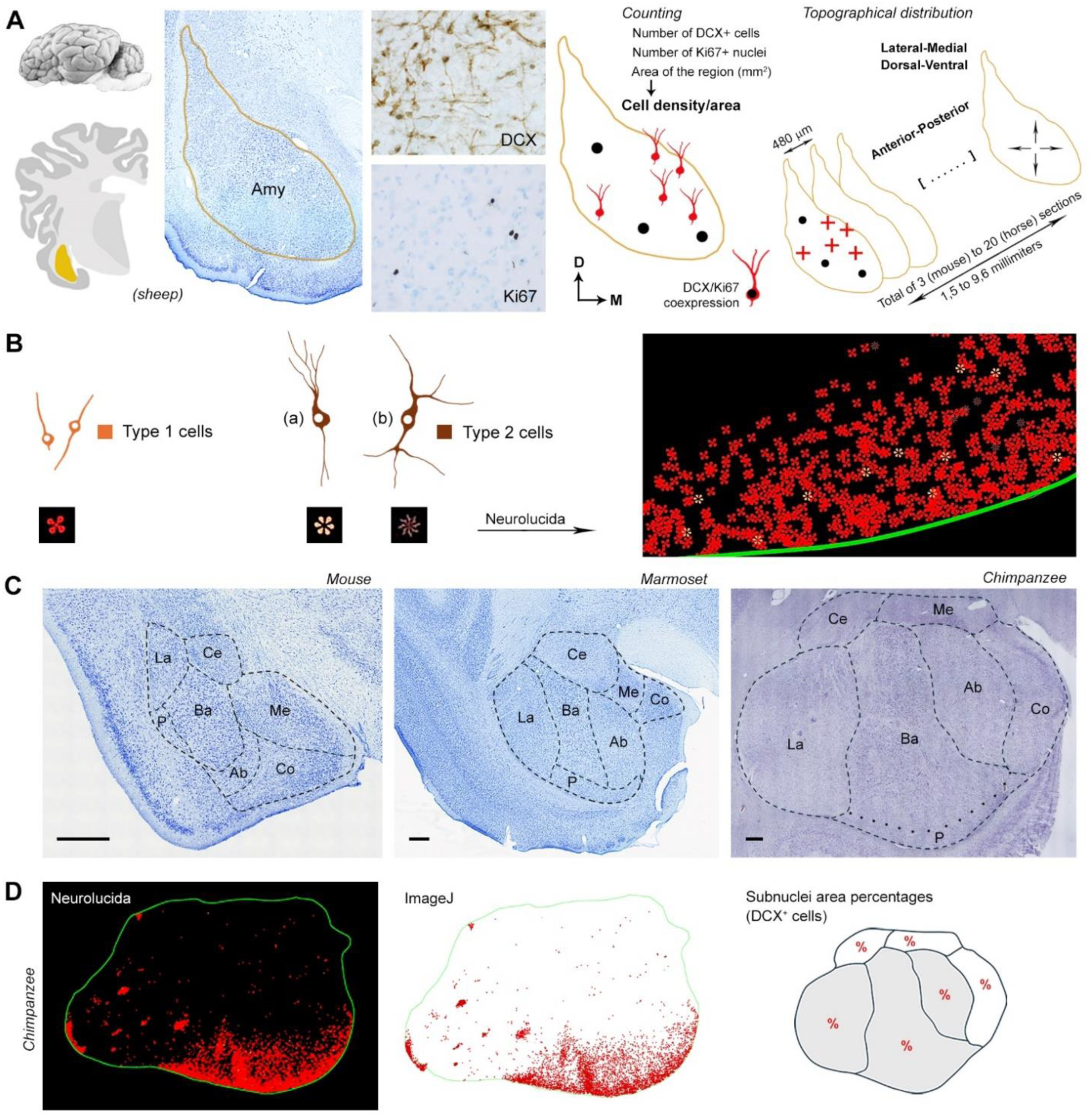
Quantitative analyses and subnuclei segmentation in the amygdala of different mammals. (**A**) Establishing amygdala areas after segmentation carried out on serial coronal sections (see Fig. 1). DCX^+^ cells (red crosses) and Ki67^+^ nuclei (dividing cells; black dots) were counted with Neurolucida software (example given in **B**) in 480 µm-spaced sections, to obtain cell density/area and cell density/volume of the entire region of interest. The anterior-posterior distribution and the bidimensional spatial distribution (dorsal-ventral and lateral-medial) of immunoreactive elements in single coronal sections was also studied (see Figs. 6 and 7). The aim was to obtain comparable cell densities and spatial distributions in all species to investigate phylogenetic variation (and age-related variation), and to check for presence/absence/amount/nature of dividing cells (see text and Table 1). (**B**) While counting the DCX^+^ cells, a distinction was made on Neurolucida between type 1, type 2a, and type 2b (the latter then considered together as type 2 cells), based on their soma size and dendritic arborization (see Fig. 2A). Colors (yellow and brown) for the two main cell types are the same in pie charts of Fig. 5C). (**C**) Amygdala subnuclei segmentation (here shown in mouse and primates; for all species and different anterior-posterior levels, see Fig. S1) was performed on coronal sections stained with toluidine blue; cresyl violet in chimpanzees) and matched with: (7, 70, mouse; 71, naked mole rat; 72, 73, rabbit; 28, 74, cat; 75, 76, sheep; 77, 78, marmoset; 46, chimpanzee; 79, horse). (**D**) To allocate the immature cells in each of the amygdala subnuclei, the Neurolucida fields employed for DCX^+^ cell counting were used; they were imported in ImageJ and the percentage of pixels highlighted in red in the selection gave the percentage of area occupied by the cells (see Table S3). Scale bar: C, 500 µm.

**Figure 5.**
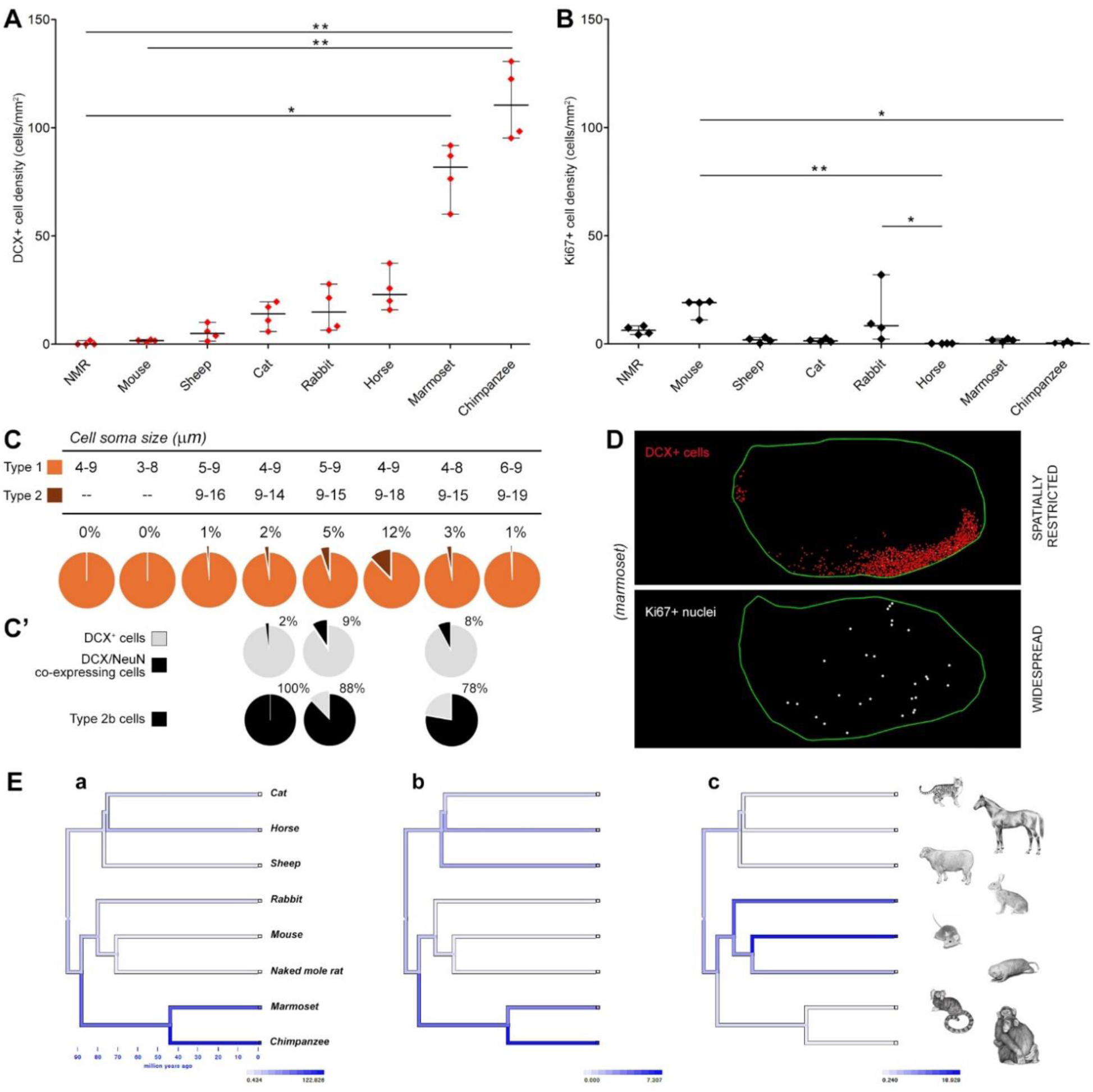
Quantification of DCX^+^ neurons and Ki67^+^ nuclei in the amygdala of young adult mammals. (**A**) Cell density and statistical analysis of DCX^+^ cells in the amygdala of eight mammalian species (listed in ascending order from left to right). A high degree of heterogeneity is detectable from rodents to primates, the latter showing the higher amount. (**B**) Cell density and statistical analysis of dividing cells in the amygdala of the same animal species and age. A rather minimal homogeneity is detectable, with slight prevalence in rodents and rabbit, and very low levels in primates (see also Fig. 6). Note the sharp contrast between DCX^+^ neuron abundance (**A**) and Ki67^+^ dividing cell scarcity (**B**) in primates. (**C**) Counting of type 1 and type 2 cells (see Fig. 2A and 4B); top, cell soma diameter ranges; bottom, percentages are showed in pie charts (numbers indicate the percentage of type 2 cells). (**C’**) Percentages of DCX^+^ neurons co-expressing NeuN (cat, rabbit, marmoset); all co-expressing elements were complex cells (type 2 cells), most of them being type 2b cells (see Fig. 3A, and data S1). A-C, Animal species are arranged from left to right according to increasing DCX^+^ cell density. (**D**) DCX^+^ neurons (mostly restricted to the ventral-medial aspect of the amygdala) and Ki67^+^ dividing cells (widespread in the entire region) have different topographical distribution, suggesting they belong to different populations (example given on marmoset; for other species see Figs. 6 and 7A). (**E**) Ancestral character state reconstructions of trait evolution for DCX^+^ cell density (a), DCX^+^ cells as a percentage of the basolateral complex of the amygdala (b), and Ki67^+^ cell density (c) mapped onto the phylogeny.

**Table 1.** Summary of the observations gathered in the present study and shared by different mammals indicating the DCX^+^ cells in the amygdala as “immature” neurons rather than newborn elements.

|  |
| --- |
| Expression and co-expression of typical markers of neuronal immaturity (DCX, PSA-NCAM) – Figs. 2 and 3A |
| Morphology reminiscent of type 1 (small soma, unipolar/bipolar) and type 2 cells (large soma, complex dendritic tree) typical of cINs (corresponding to different maturational stages; Rotheneichner et al., 2018; La Rosa et al., 2020) – Figs. 2A |
| Complex (type 2b) cells reminiscent of the principal neuronal type in amygdala – Fig. 2A |
| High percentage of type 1 cells and low percentage of type 2 cells - Fig. 5C |
| Low percentage of DCX<sup>+</sup> cells (mostly type 2, complex cells) co-expressing the maturational marker NeuN (Fig. 5C') |
| Different topographical distribution of DCX<sup>+</sup> cells with respect to dividing cells (the former grouped in restricted areas and constantly associated with specific subnuclei; the latter widespread through the entire amygdala), indicating they belong to distinct populations - Figs. 5 and 6 |
| Absence of co-expression with Ki67<sup>+</sup> nuclei, indicating they are not dividing - Fig. 3B' |
| High-rate co-expression between Ki67<sup>+</sup> nuclei and glial markers (Olig2, SOX10), indicating that most dividing cells in the amygdala are glial elements - Fig. 3C |
| Frequent detection of Ki67<sup>+</sup> “duplets” indicating OPC division - Fig. 3C |
| Remarkable variation in number (cell density) between mammalian species, as previously described for cINs (La Rosa et al., 2020) - Fig. 5A |
| Substantial maintenance of the IN cell populations at adult and old stages, in non-rodent species - Fig. 8 |

Overall, the DCX^+^ cell types of the amygdala were reminiscent of those described in the cortex (6, 8, 12), thus suggesting that populations of immature neurons characterized by similar, progressive stages of maturation are likely shared by cerebral cortex and amygdala, their final product being tailored to the specific neuronal types of the two regions. These features prompted us to identify these cells to be neurons in arrested maturation (INs). On this basis, we sought to determine whether some of the DCX^+^ cells in the amygdala might be actively dividing, by performing DCX/Ki67 antigen double staining for confocal analysis to search for possible marker co-expression in all the species considered (Fig. 3B’). After counting a total of 856 Ki67^+^ nuclei in 252 confocal fields belonging to all species studied (aside from chimpanzees, not investigated in confocal microscopy), no co-expressions with DCX^+^ cells was ever detected, indicating that the DCX^+^ immature neurons and the proliferating cells in the amygdala most likely belong to distinct cell populations, as previously found in the cerebral cortex (15). Based on this result, and on the frequent appearance of the Ki67^+^ nuclei as “doublets” displaying the typical characteristic of dividing oligodendrocyte progenitor cells (OPCs; 40; Fig. 3C), the possible co-expression of the cell division marker with glial markers was investigated. Although NG2 (Neuron Glia Antigen 2) is considered the ideal molecular marker for the identification of early developing OPCs in rodents, its expression cannot be easily detected in postmortem tissues, which are usually too heavily fixed (41). Accordingly, the marker worked well in perfused mice brains (see 42) while giving poor results in most other mammalian brain tissues. As an alternative, we employed the pan-oligodendrocyte transcription factors SOX10 (43) and Olig2 (44) in double staining experiment against Ki67 antigen. Indeed, co-expressing cells were frequently observed in all the specimens examined (Fig. 3C), thus confirming the glial nature of most dividing cells in the amygdala.

### Quantification of DCX^+^ neurons and dividing cells (cell densities) in the amygdala of different mammals

The number of DCX^+^ cells per mm^2^ of amygdala was assessed through measurements on Neurolucida software in serial coronal sections spaced 480 µm apart, by counting all the cells in the amygdala segmented area (cell density/area; Fig. 4A,B). Due to the different length of the structure in each animal species (from 1,4 mm in rodents to 9,6 mm in horses), different numbers of sections were considered (from 3 to 20; Fig. S1 and Table S4). This approach allowed us to obtain a comparable value in all species, whatever the size of amygdala and the spatial arrangement of the DCX^+^ cells within the region. Given the non-homogeneous distribution of the latter (see below), a stereological method for quantification was excluded in favor of a direct cell counting, also to make the analysis as close as possible (comparable) to that previously performed in the cerebral cortex (15).

In Fig. 5A are reported the results for quantifications in young adult animals, from all the species in this study (indicated by the green line in Fig. 1). All non-rodent species (marmoset, rabbit, cat, sheep, chimpanzee, horse) showed a higher density of DCX^+^ cells with respect to rodents (mouse, naked mole rat). Both primates (marmoset, chimpanzee) had significantly greater density of DCX^+^ cells (nonparametric Kruskal-Wallis test, p<0.05), with a two orders of magnitude difference between mouse and chimpanzee (Fig. 5A; tabulated data can be found in data S1). Differential counting of type 1 and type 2 cells was also performed (percentages reported in pie charts of Fig. 5C). The small, highly immature cells were far more numerous (usually exceeding 90%) than the large, ramified cells. The same analysis was carried out on Ki67^+^ nuclei. In comparison with data obtained for the DCX^+^ cell population, the density of Ki67^+^ dividing cells showed a quite different pattern, generally at low densities in all species (Fig. 5B). Within this range, the rate of cell division was higher in rodents and more sporadic in non-rodent species, with a significant difference in mouse with respect to chimpanzee (nonparametric Kruskal-Wallis test, p<0.05; Fig. 5B).

The reconstruction of evolutionary change in the values of DCX^+^ cell density, DCX^+^ cells as a percentage of the basolateral complex of the amygdala, and Ki67^+^ cell density is shown in Fig. 5E. Increased density of DCX^+^ cells occurred along the primate lineage, whereas an increase of Ki67^+^ cells occurred in a separate branch of the phylogeny that includes rodents and rabbits.

Overall, rodents displayed low DCX^+^ cell density and higher Ki67^+^ cell proliferation rate, while this pattern was inversed in primates, a general pattern further supporting the idea that INs and cell divisions in amygdala represent different cell populations. Also, the topographical distribution of the Ki67^+^ nuclei and DCX^+^ cells appeared quite different in all species considered, with the former randomly distributed in the entire coronal plan and the latter mostly grouped in the ventral portion of the amygdala (Fig. 5D), prompting further investigation of the spatial distribution of the two cell populations.

### Topographical distribution of DCX^+^ neurons and Ki67^+^ dividing cells in the amygdala of mammals

Most DCX^+^ cells appeared as isolated and rare elements in rodents (mouse, naked mole rat), while frequently organized to form clusters and complex networks in other species (Fig. 2B). The topographical distribution of the cells in the region was analyzed both in the entire longitudinal, anterior-to-posterior axis (Fig. 6 and Fig. S1) and in the coronal plan (medial-to-lateral/ventral-to-dorsal axes; Fig. 7), in all species considered. The anterior-to-posterior distribution was obtained after generating histograms of the total cell number for each of the coronal sections considered (see Methods; Figs. 6 and Fig. S1). This showed remarkable interspecies variation, with the DCX^+^ cells concentrated in different amygdala compartments in different species (e.g., mostly posterior in cat and sheep, while mostly in the middle in horse and primates).

**Figure 6.**
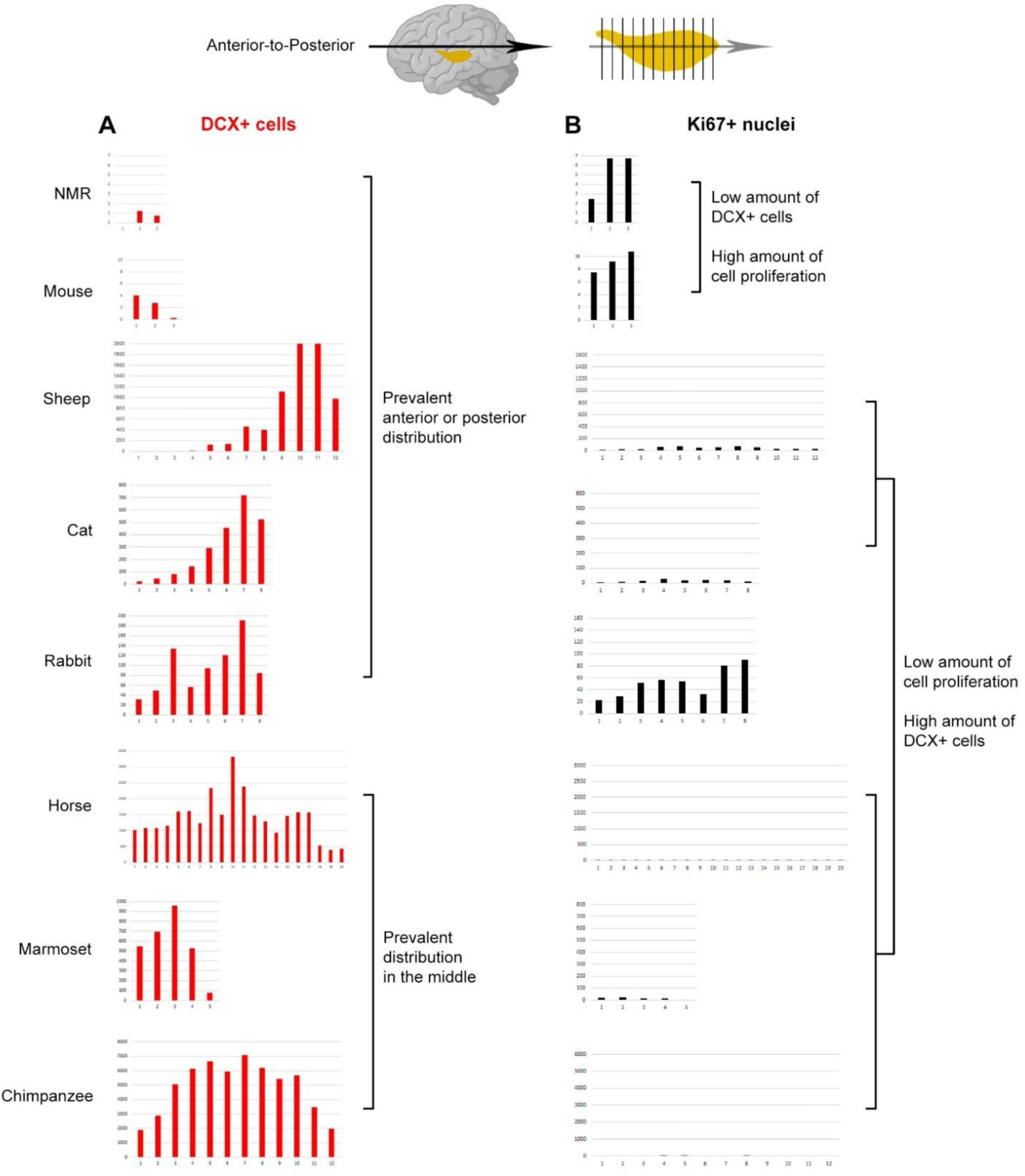
Spatial distribution in the amygdala anterior-posterior (longitudinal) axis. The amount of DCX^+^ cells (density/area; red) and Ki67^+^ nuclei (black) are reported as histograms for each single coronal section of the amygdala counted in each species (see Fig. S1), in the sequence from the anterior (left) to posterior part (right), thus revealing the pattern of longitudinal spatial distribution of these cell populations. While the distribution of the DCX^+^ cells was highly heterogeneous among mammals, e.g., far higher in the anterior (mouse), middle (primates) or posterior part (sheep, cat), that of dividing cells was quite homogeneous, especially in gyrencephalic species and marmoset. Animal species are arranged from top to bottom according to increasing DCX^+^ cell density.

**Figure 7.**
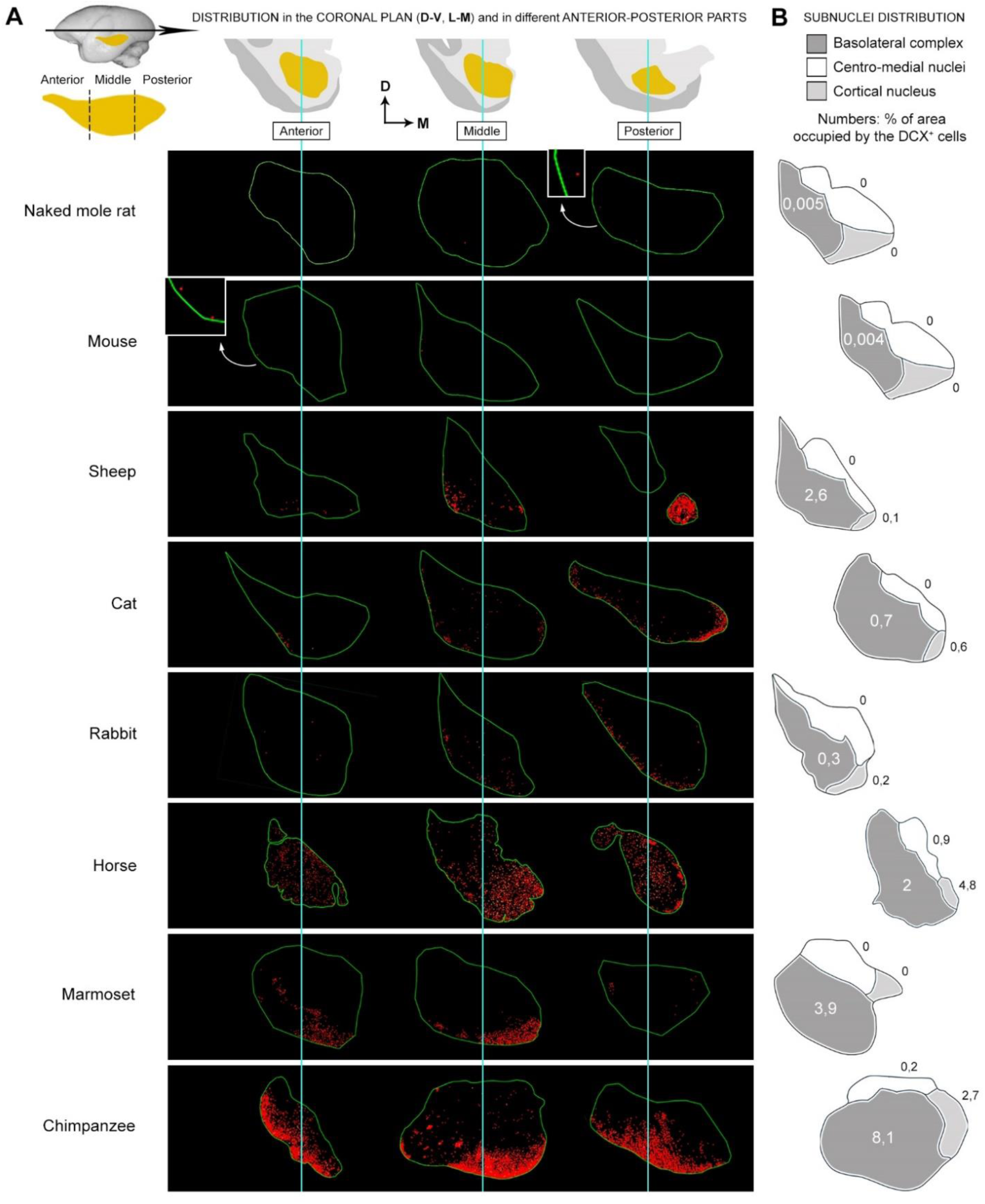
Spatial distribution in the amygdala coronal plan. (**A**) Topographical distribution of DCX^+^ cells within the amygdala of different mammals visualized in Neurolucida fields obtained from brain coronal sections used for cell counting (red dots: DCX^+^ cells; green line: amygdala perimeter). Top, the anterior-to-posterior extension of the amygdala has been split in three parts (anterior, middle, posterior; see also Fig. S1) to compare the distribution in different coronal plans with that observed in the longitudinal axis. While the DCX^+^ cell distribution in the anterior-to-posterior extension of the amygdala appeared to be highly heterogeneous among species (Fig. 6), that in the coronal plan was quite constant, being prevalent in its ventral part. (**B**) Percentages of areas occupied by the DCX^+^ cells within the three main amygdala subdivisions (BLc, centro-medial complex, and cortical nucleus), indicating the prevalent association of the INs with nuclei of the basolateral complex (BLc, dark grey), and, to a lesser extent, with the cortical nucleus in large-brained species (horse and chimpanzee; light grey). The invasion of the BLc is particularly evident in primates, sheep, and horse; in addition, despite the representation of amygdala and subnuclei is not in scale, the relative volume of the BLc is far greater in primates than in rodents (45, 48), thus making the percentages of areas occupied by the INs even higher. The very small percentages of areas occupied by the INs in rodents do correspond to their highly restricted location within the small paralaminar nucleus, without invading the BLc. Percentages for each of the subnuclei are reported in Table S3. Animal species are arranged from top to bottom according to increasing DCX^+^ cell density.

By contrast, the distribution of the cells in the coronal plan was consistent across species, being prevalent in the ventral aspect of the amygdala (Fig. 7A), thus indicating that it might be a conserved trait, with the immature cells being mainly associated with specific subnuclei. To examine this further, the different amygdala subnuclei were identified by combining our anatomical mapping with existing atlases and previous descriptions in each species (see Fig. 4C and Fig. S1 for detail). Due to heterogeneous knowledge and terminology existing in the literature concerning the amygdala subnuclei across mammals (37, 45–48), we adopted a simplified grouping which considers the standard subdivisions of the basolateral complex (lateral, basal, and accessory basal nuclei), the centro-medial complex (medial and central nuclei), and the cortical nucleus (Fig. S1). In previous reports of comparative neuroanatomy, the paralaminar nucleus has not been identified and characterized as a separate nucleus in all species, being often considered as part of the basal nucleus (47, 49); for this reason, and due to its cellular composition (see Discussion), it has been included in the basolateral complex, referred hereafter to as BLc.

We found that most DCX^+^ neurons were concentrated in the area of the BLc. To allocate more precisely the immature cells to each of the amygdala subnuclei, the percentage of area occupied by the cells were measured (see Methods and Table S3). As shown in Fig. 7B, most of the DCX^+^ cells were consistently located in the nuclei forming the BLc of each species (apart from rodents, in which a few INs are segregated in the small paralaminar nucleus; see also Ref. 7). Notably, the association of the INs with the BLc was independent from variation in their density or anterior-posterior distribution, again suggesting that it is a conserved trait. The percentage of area occupied by the INs in the BLc varied across the species, with the greatest amount in primates, in parallel with DCX^+^ cell density (Table S3 and Fig. 7B).

Overall, these data demonstrate phylogenetic variation in amygdala INs, with a focus of species differences in the BLc.

No DCX^+^ cells, or only a negligible amount, were found in the medial and central nuclei of the amygdala in both rodent and non-rodent species, while a modest number of them were present in the cortical nucleus in horses and chimpanzees (Table S3).

As showed above (Fig. 5D), by comparing Ki67-stained and DCX-stained adjacent sections the difference in the distribution of the two cell populations was evident, the former being randomly scattered in the entire amygdala while the latter mostly grouped in its ventral part (Fig. 5D). This difference was even more evident quantitatively, both considering whole cell density medians (Fig. 5A,B) and those representing single coronal sections (comparison between left and right histograms in Fig. 6). Hence, quantitative analyses combined with topographical location of the cells confirm that INs and dividing cells in the amygdala belong to distinct populations.

### Quantification of DCX^+^ cells, dividing cells, and amygdala volumes at different ages

To investigate whether the amount of DCX^+^ and dividing cells change during the lifespan, we analysed brains from the young adult stage to other available ages, including middle age in mouse, naked mole rat, marmoset, cat, sheep, horse, and old age in mouse and chimpanzees (Figs. 1 and 8). A prepuberal stage was also evaluated in mouse, naked mole rat, rabbit, and sheep.

**Figure 8.**
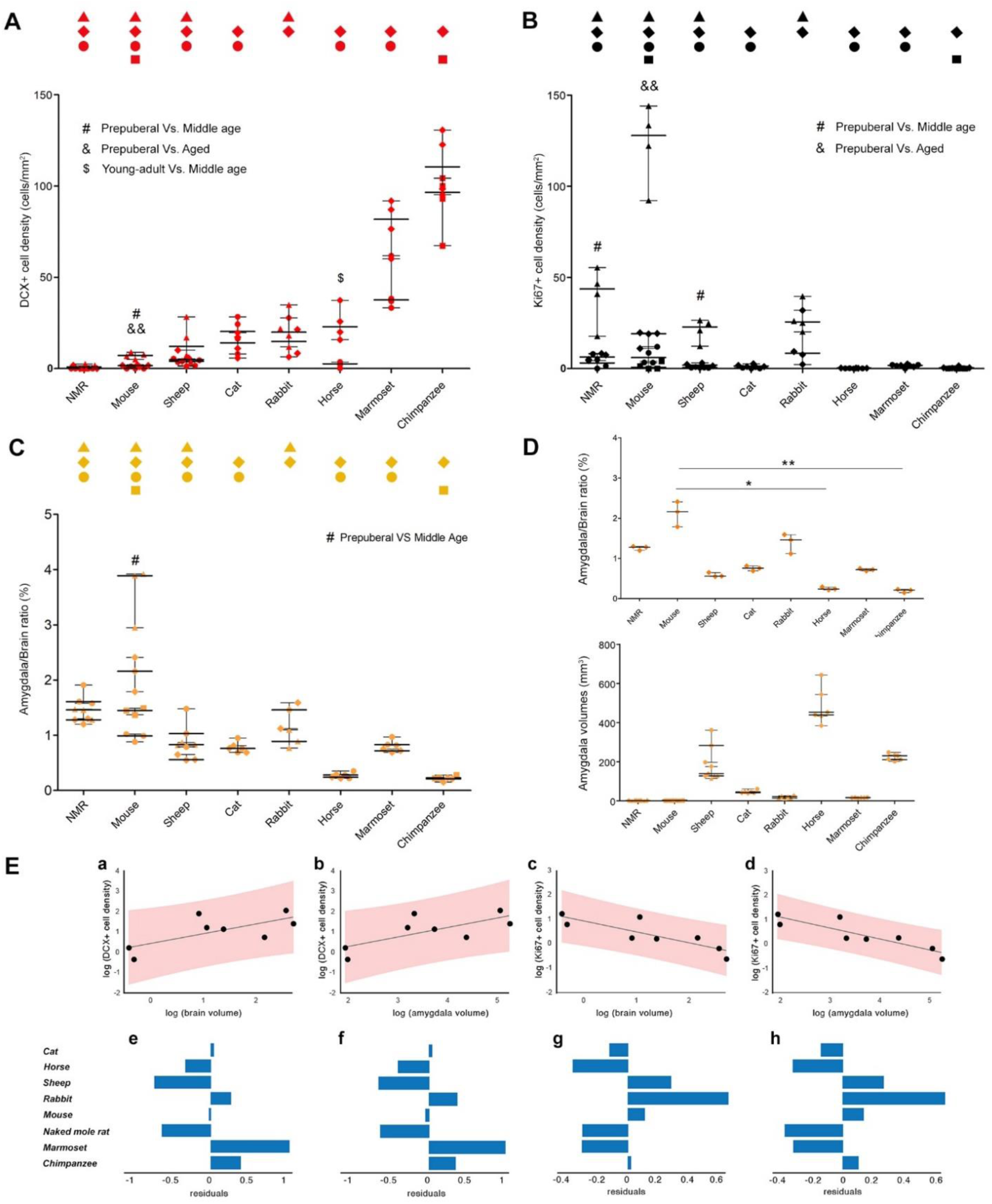
Quantification of DCX^+^ neurons, Ki67^+^ nuclei, and amygdala volumes at different ages. (**A**) Cell density and statistical analysis of DCX^+^ cells in the amygdala of eight mammalian species at different ages (all ages investigated are included; symbols indicating age groups are indicated in Fig. 1A). (**B**) Density and statistical analysis of dividing nuclei in the amygdala of the same animal species and ages (all ages investigated are included). (**C**,**D**) Amygdala volume estimation. (**C**) Amygdala/brain volume ratio at all ages investigated; note the substantial invariance of volumetric ratio through different animal species and ages. (**D**) Amygdala/brain volume ratio at young adult age (top), and amygdala real volumes in each animal species at different ages (bottom); while the absolute volume of the subcortical region is higher in large-brained species (as expected), the volumetric ratio with respect to the whole brain is substantially stable, slightly higher in small-brained ones. A-D, Animal species are arranged from left to right according to increasing DCX^+^ cell density. (**E**) Least squares regression of DCX^+^ cell density against brain volume (a) and amygdala volume (b), with associated residuals of the regressions shown underneath (e and f). Least squares regression of Ki67^+^ cell density against brain volume (c) and amygdala volume (d), with associated residuals of the regressions shown underneath (g and h). All regression plots are on a log scale and show the 95% prediction intervals.

A substantial drop (statistically significant in mice) was observed by comparing prepuberal with young adult stage (Fig. 8). This fact is consistent with previous reports indicating that INs, as other forms of plasticity, mainly characterize juvenile ages, both in amygdala (7, 30) and cerebral cortex (15). Nevertheless, in large-brained gyrencephalic mammals (apart from horse) and primates, the age-related changes were not statistically significant, indicating a substantial stabilization in adulthood (Fig. 8; nonparametric Mann-Whitney test, p<0.05). In a previous report carried out in the mouse amygdala, the DCX^+^ neurons have also been reported to drop in number very rapidly within the first two months of life (7), and we confirm here that only a few cells remain thereafter, being negligible at middle age and old stages (0,25 and 0,08 cells/section, respectively; see Table S5 and Fig. 9). Even in another rodent species characterized by extended lifespan and retention of neotenic features, such as naked mole rats (50), no more DCX^+^ cells were detectable from middle age onward (Table S5). By contrast, in primates, particularly chimpanzees, the amount of scINs appeared rather unchanged, even between relatively distant life stages (e.g., young adult and old; Figs. 8 and 9). By comparing mice and chimpanzees at these correspondent life stages, the DCX^+^ neurons (total number estimation/hemisphere; see Table S5) dropped from around 70 in young adults to 3 in aged stages (24-fold reduction) in the former, and from 700,000 to 470,000 (only half reduction) in the latter.

**Figure 9.**
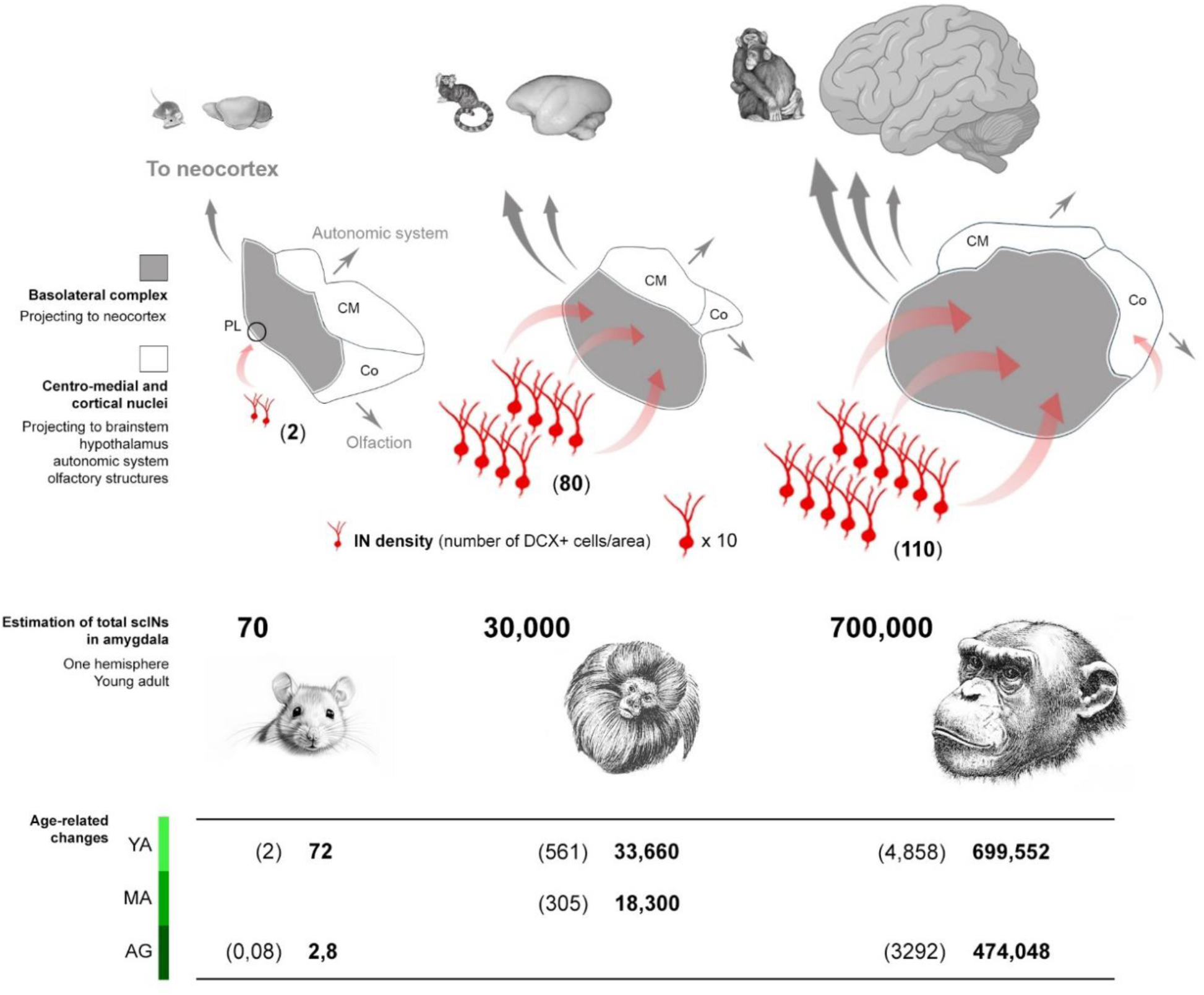
Occurrence of DCX^+^ immature neurons in the amygdala of mouse and primates: relationship with evolution of subnuclei scaling and cortical connectomics. Top and middle: the mammalian amygdala can be split into two main parts considering the basolateral complex (BLc, grey; here considered including the paralaminar nucleus), the centro-medial nucleus (CM), and the cortical nucleus (Co). While CM and Co do not change a lot in their relative volume among animal species, the BLc is remarkably larger in primates (up to 60-70%) with respect to rodents (around 30%; here not visible since not in scale and because linked to the entire anterior-to-posterior volume of the region); such an expansion has been related to increasing projections to the neocortex typical of primates (grey arrows; 45,48; see text). Despite a huge difference in the number of INs (red cells) found in primates with respect to rodents, as well as differences in the anterior-posterior distribution, their prevalent occurrence within the basolateral complex is a maintained trait (some INs were found in the cortical nucleus of horse and chimpanzee; not represented here, see Fig. 7B). While mostly restricted to a very small paralaminar nucleus (PL) in rodents, the scINs largely invade the basolateral nuclei in primates (red arrows and Fig. 7B). After scIN total amount estimation in the amygdala of each hemisphere (bottom), chimpanzees possess a reservoir of undifferentiated cells four orders of magnitude higher than mice, and this reservoir is maintained through ages in primates, while decreasing in rodents (quantitative data extracted from Table S5). In brackets, average number of DCX^+^ cells in a coronal section of the amygdala; in bold, estimation of total number of DCX^+^ cells/hemisphere. YA, young adult; MA, middle age; AG, aged.

In parallel with the analysis of DCX^+^ cell density, counting of Ki67^+^ nuclei was also performed in all species and ages available. Again, a significant drop was observed in species analyzed at prepuberal stages (Fig. 8B). Particularly in rodents, which showed higher rates of cell division than other species, a significant drop occurs at juvenile stages. Subsequently, the levels of cell proliferation do not change significantly at later stages, remaining low in adulthood in rodents and non-rodent species (nonparametric Mann-Whitney test, p<0.05; Fig. 8B). Interestingly, rodents and primates again substantially differ from each other, the former having a higher amount of parenchymal cell division (still present at young adult stages) while the latter, along with other large-brained gyrencephalic species, displayed very low levels at all life stages (Fig. 8B).

Finally, an analysis of amygdala volumes in different species at the same age (young adult stage) and at different ages was carried out, to measure both absolute volumes and amygdala/brain volume ratio (Fig. 8C,D; tabulated data can be found in data S1). While the absolute amygdala volume, as expected, changed considerably with increasing brain size (from 1,9 mm^3^ in mouse to 460 mm^3^ in horse; Fig. 8D, bottom), the amygdala/brain volume ratio was substantially similar in all species, thus independent from brain size (Fig. 8D, top).

To explore the scaling relationship between DCX^+^ and Ki67^+^ cell densities with brain and amygdala volumes, least squares regression analyses were conducted (Fig. 8E). For the relationship between DCX^+^ cell densities with brain volume and amygdala volume, the results indicated a significant positive association (brain volume: y = 0.419 + 0.479x; R^2^ = 0.50, F(1,6) = 5.93, p = 0.051; amygdala volume: y = −0.661 + 0.467x; R^2^ = 0.52, F(1,6) = 6.42, p = 0.044.) In contrast, the relationship between Ki67^+^ cell density with brain volume and amygdala volume displayed a significant negative association (brain volume: y = 0.950 - 0.448x; R^2^ = 0.72, F(1,6) = 15.24, p = 0.008; amygdala volume: y = 2.003 - 0.448x; R^2^ = 0.79, F(1,6) = 22.13, p = 0.003).

## Discussion

### DCX^+^ cells in the amygdala as neurons in arrested maturation

In the present study, at least ten features commonly attributed to neurons in arrested maturation or “immature neurons” (INs) previously described in the cerebral cortex (1, 2, 5, 8) were consistently associated with the DCX^+^ neurons residing in the amygdala of widely different mammals, indicating the existence of scINs rather than newborn neurons (Table 1). In summary, these cells strictly fit with the morphological subtypes described for the progressively maturing neurons of cortical layer II, with a greater prevalence of small, simple-shaped cells and a minor fraction of complex morphologies reminiscent of the principal cell type of the region (pyramidal neurons in the cortex, Class I neurons in the amygdala). Most of the scINs co-express the markers of immaturity DCX and PSA-NCAM, while a small subpopulation of large and complex cells (type 2 cells, mostly type 2b) also express NeuN, an RNA-binding protein detectable in postmitotic neurons that have begun differentiation (51); the scINs never co-express nuclear proteins associated with cell division, and display an utterly different spatial distribution with respect to proliferating cells, which are mostly identifiable with parenchymal glial elements known to represent the larger population of proliferative cells in the brain (40, 52, 53). Also, the remarkable phylogenetic variation observed in the amygdala, with a large prevalence of scINs in primates compared to rodents (with an inversion of DCX^+^ cell and Ki67^+^ dividing cell densities in the two groups of animals) points to their nature as neurons in arrested maturation.

The existence of scINs is further supported by previous reports carried out in sheep and mice injected as embryos with 5′-bromo-2′-deoxyuridine, in which the DCX^+^ cells in the amygdala and claustrum of lambs (29) and those in the amygdala of mouse pups (7) were found to be prenatally generated, similar to the cINs. Though these experiments cannot be easily replicated in most large-brained, gyrencephalic species, the convergence of multiple observations strongly indicates that populations of scINs do exist in the amygdala of mammals. Overall, multiple lines of experimental evidence converge to define these cells as a reservoir of non-dividing, young neuronal elements independent from stem cell-driven neurogenesis and reminiscent of their counterpart in the cortex (15).

### Interspecies variation in the amygdala of mammals

In the present study, densities of DCX^+^ scINs were evaluated in a comparable way within the amygdala of phylogenetically and neuroanatomically diverse mammals by using the same method across the 80 brains analyzed and following the approach previously employed to study the phylogenetic variation of cINs (15). As a result, remarkable interspecies differences were found: the scINs were extremely rare in rodents and more abundant in large-brained gyrencephalic species, including perissodactyls, artiodactyls, carnivorans, and primates (Fig. 5A). The density of scINs was also positively correlated with brain and amygdala volume (Fig. 8E). This general pattern was reminiscent of that reported for cINs, whose densities were also related to brain size and gyrencephaly (15). Additionally, beyond allometric scaling, the scINs seem to be more numerous especially in primates (which in our sample included both large-brained, gyrified chimpanzees and small-brained, mostly lissencepahlic marmoset monkeys). In contrast, few cINs were observed in the neocortex in marmosets (15). Future studies should examine a greater range of primate species that vary in brain size to confirm that the observed increase in amygdala scINs is consistent in this phylogenetic group. In the cerebral cortex, such variation has been recently explained as a trade-off between different types of developmental processes involving stem cell-driven adult neurogenesis versus neurons in arrested maturation (17). This might have occurred in brain evolution of mammals as a mechanism to assure the most appropriate type of plasticity is available, tailored for the brain size, life history characteristics, and behavioral adaptations in each species. Following this view, rodents have maintained a dependence on olfaction to navigate in their environments, while larger-brained species show specializations for experience-dependent learning related to neocortical expansion (54). On this basis, the high number and widespread occurrence of INs found in the amygdala of primates might be linked to the increasing importance of this subcortical region in the integration of complex social interactions that can be crucial for their survival and reproductive success (26, 48, 49, 55). Different groups of primates, while widely differing in their neocortical size, maintain a rather constant amygdala/brain scaling among them and with other mammals (56–58). Primates share an increasing importance of amygdala circuits/functions with respect to rodents (48, 49), including their connections with the cortex (45), a fact that is reflected in amygdala subnuclei scaling (discussed below).

### Phylogenetic variation of amygdala matches subnuclei scaling

While the general amygdala/brain volume ratio has remained relatively unchanged among widely different mammals (56-58; confirmed here by volume estimation in all species, see Fig. 8C,D), some features of amygdala anatomy display phylogenetic variation, specifically in the scaling patterns of individual nuclei (45, 46, 48, 49, 58). The lateral, basal, and accessory basal nuclei are proportionally larger in primates than in rodents, with respect to other nuclei (central, medial, and cortical) which differ relatively little (45, 46, 48, 49). These changes are also reflected in connectivity: the amygdala basolateral nuclei are broadly connected to the neocortex, while the centro-medial and cortical nuclei mostly communicate with brainstem, hypothalamus, autonomic system, and olfactory structures (45, 48, 59). Accordingly, the increase in the basolateral complex volume has been correlated with increases of neocortical volume among primates, as a response to increased processing demands from the neocortex (56). In line with the expansion of the associated cortical territories, the subnuclei of the BLc are preferentially expanded in primates compared with rodents (62% of total amygdala volume in macaques, and 69% in humans, with respect to 28% in rats; 45, 48), and, compared to rodents, the primate amygdala has lower neuronal density and larger neuropil volume, which have been associated with greater complexity in dendritic arborizations and axonal innervation, enabling a greater capacity to integrate information (45). Hence, in contrast to the cortico-medial region which is linked with olfaction and automatic, defensive reactions, the BLc has predominant connections with the neocortex (48, 49; Fig. 9).

Here we show that the remarkable increase in the number of scINs between rodents and primates (with other species intermediate), appears to be associated most closely with the BLc, following its phylogenetic expansion (Fig. 9). Even though the proportional size of the whole amygdala does not differ across the species considered, the phylogenetic variation of the scINs matches with its expanding subnuclei and connectome, similarly to what has been observed for cINs in the expanded neocortex of gyrencephalic species (15). Of importance, we show that in large-brained gyrencephalic species and primates, the occurrence of immature cells goes well beyond the paralaminar nucleus, an area that in most mammals is not classified as a subregion in its own, likely because it appears mostly composed of a mass of small, densely packed immature cells rather than having a specific histology. Although the fundamental function of the amygdala in fear and emotional learning is conserved across species, the region is under greater influence of cortical activity in primates, integrating contextual information linked to more complex behaviors such as social interaction and cognition (45, 48, 49). Due to its extensive connections with much of the cortex, the amygdala is thought to be involved in the processing of salience, significance, ambiguity, and unpredictability, thus playing a role in selectively processing the inputs that are the most relevant to the goals of the animal (60). Considering that the regions of high expansion mature later in development to take advantage of experience and social learning (61), the greater occurrence of scINs in this region may provide a substrate for plasticity in the form of a reservoir of undifferentiated cells subserving highly sophisticated functions.

### A reservoir of subcortical INs is maintained through the lifespan in primates

In addition to interspecies analysis, the amount of DCX^+^ neurons and proliferating cells was investigated longitudinally in each species to examine possible age-related variation (Fig. 1A). Results and statistical analysis are shown in Fig. 8A,B. For DCX^+^ cells (Fig. 8A), despite a progressive age-related reduction observed in all species (significant in mouse and horse), the number of INs was rather stable in adult primates. This pattern is particularly evident when comparing mice and chimpanzees, both studied at young adult and aged stages (Fig. 8A and 9): the chimpanzees, besides having far more abundant scINs, maintains many of them through adult and old ages, while they appear to be depleted very early in mice. Our results in mice are consistent with a recent report wherein DCX^+^ cells were investigated in the amygdala of young animals (P7 to P60; 7), showing a sharp decline through adolescence and reaching very low levels in the 2-month-olds. We extended the analysis to older ages and found only negligible numbers of DCX^+^ cells (0,25 and 0,08 cells/section in middle age and aged mice, with only 3 total cells in the entire amygdala of aged animals; Table S5). By contrast, estimations of the total number of scINs per hemisphere in chimpanzees revealed a reservoir of 470,000 cells in aged individuals, which remained from the nearly 700,000 of young adult animals (Table S5 and Fig. 9), suggesting that these undifferentiated cells may play a substantial role (at present still unknown) through the lifespan. In accord with this view, confocal analyses to examine co-expression of DCX and PSA-NCAM indicated that most scINs remain in a state of immaturity at adult and old ages, while the percentage of those displaying a sign of starting maturation (co-expression of NeuN) was very low (around 2-9% of all DCX^+^ cells, all corresponding to type 2, “complex” cells, mostly type 2b cells; Figs. 3A and 5C’). Overall, the age-related trend of scINs is reminiscent of that described in the cerebral cortex, where the cINs also undergo a dramatic drop at young ages in mice (similarly to adult neurogenesis; 42) while remaining at higher levels in large-brained gyrencephalic species, especially chimpanzees (15, 42). Yet, a different ratio can be observed between type 1 and 2 cells in cortex and amygdala: the reservoir of highly immature cells (type 1 cells) is larger in amygdala (the type 2 cells being restricted to a range from 0 to 2% - except for horses; Fig. 5C) than in the cortex (the cINs showing a range of 3-14 %, reaching 44% in mice; see 15). Accordingly, in a descriptive study performed on the human cerebral cortex at different ages spanning from neonatal to very old stages, many DCX^+^ neurons of layer II (cINs) were still present in the neocortex of old individuals while the occurrence of DCX^+^ cells in the hippocampus of the same specimens were quite reduced (20).

Overall, our approach across mammalian species and ages reveals that in non-rodent species (especially primates, and mostly long-living species) the amygdala may rely on abundant populations of highly undifferentiated scINs, whose amount does not deplete sharply in juvenile ages, reaching a stabilization in adult and old individuals. A previous study performed in humans by Sorrells et al. (30) mainly focused on the drop in the amount of DCX^+^ cells after adolescence. They also described the persistence of immature cells in adult and old individuals (up to 77 years of age), though their analysis was performed on a few adult/old human samples using tissue blocks from the temporal lobe, in which the amygdala was not considered in its entire extension in the coronal plan, their quantitative analysis being restricted to the paralaminar nucleus. The findings of our systematic study in two primate species, considering the whole volumetric extension of the amygdala, agree with the Sorrells study in most respects, yet suggests that the human amygdala may host more scINs than currently thought.

### Concluding remarks

We showed here that a population of young neurons (scINs) sharing features with those previously described in the cerebral cortex (cINs) is present in the amygdala of widely different mammals, displaying remarkable phylogenetic variation. Their density spans nearly two orders of magnitude from mouse to chimpanzee, and an estimation of their total amount in each hemisphere indicated a four-order magnitude difference, with 700,000 scINs in the 5,7 mm long structure of chimpanzee amygdala (Fig. 9 and Table S5). By considering these numbers, together with their counterpart in the cortex (the cINs were estimated around 2 million cells/hemisphere in the chimpanzee; 15), the cINs and scINs are candidates for the most consistent reservoir of immature/plastic cells in primate brains, thus representing the only example of a substrate for structural plasticity in mammals that increases in association with brain size and complexity.

Overall, the scINs might represent a reserve of young cells tailored for increased complex cognitive functions in highly social mammals, supporting the idea that they may have been selected across evolution (along with the cINs) in a trade-off with stem-cell driven neurogenesis, which is more active in short-living animal species mostly relying on olfaction for their survival (e.g., rodents; 16, 17). Notably, the current findings link the scINs to the expanding subnuclei of the amygdala’s BLc, which are highly interconnected with the cerebral cortex, thus underlying the importance of cells in arrested maturation in the amygdala-cortex interaction. Since emotion and social dysfunctions arise in psychiatric illnesses through altered connectivity between the amygdala and cortical regions (62), the reservoir of young neurons in arrested maturation existing in primate brains may have future developments in the prevention and therapy of neurodevelopmental disorders, possibly as a broad substrate for the brain reserve (63, 64). These advances in our knowledge can provide the correct translation of research results obtained from different animal models, particularly laboratory rodents.

## Materials and Methods

### Brain tissues and legal permission

Brains of the different animal species used in this study were collected from various institutions and tissue banks, provided with the necessary authorizations (Table S1). All experiments were conducted in accordance with current laws regulating experimentation in each country/institution providing the brain tissues (see below description of each group of animals).

*Mus musculus* (Mouse) and *Oryctolagus cuniculus* (Rabbit) brains came from the Neuroscience Institute Cavalieri Ottolenghi (NICO) animal facility (Orbassano, Turin; Italy). *Heterocephalus glaber* (Naked mole rat; NMR) from the School of Biological & Chemical Sciences, Queen Mary University of London, London. *Callithrix jacchus* (Marmoset) brains were provided by the University of Zurich, Switzerland and from the Department of Anthropology, The George Washington University, Washington DC, USA. *Felis catus domestica* (Cat) and *Equus caballus* (Horse) brains from the Mediterranean Marine Mammal Tissue Bank (MMMTB) of the University of Padova, Department of Comparative Biomedicine and Food Science, Italy. *Ovis aries* (Sheep) both from the MMMTB of the University of Padova, Italy, and from the INRA research center, Nouzilly, France. *Pan troglodytes* (Chimpanzee) come from the Department of Anthropology, The George Washington University, Washington DC, USA.

Four prepuberal mouse (*M. musculus*; C57BL/6 mice raised at the NICO facility, courtesy of Serena Bovetti and Charles River Laboratories, RRID:MGI:3696370) brains were extracted a few minutes following euthanasia in accordance with Schedule 1 of the Animals (Scientific Procedures) Act 1986, and fixed by immersion. Four young adult, four middle-aged and four aged mouse were used. For these brains, transcardiac perfusion was performed under anesthesia (i.p. injection of a mixture of ketamine, 100 mg/kg, Ketavet, Bayern, Leverkusen, Germany; xylazine, 5 mg/kg; Rompun) with 4% paraformaldehyde in 0.1 M sodium phosphate buffer, pH 7.4. Brains were postfixed for 4 hours.

Four prepuberal naked mole rat (*H. glaber*) brains were extracted a few minutes following euthanasia in accordance with Schedule 1 of the Animals (Scientific Procedures) Act 1986, and fixed by immersion, while four young adult and four middle-aged animals were intracardially perfused with 4% paraformaldehyde in 0.1 M sodium phosphate buffer, pH 7.4 (Dr. Chris G. Faulkes, London, UK). Brains were then postfixed overnight.

Three adult marmosets (*C. jacchus*) were obtained from the Veterinary service of the University of Zurich. Animal handling and tissue collection was conducted according to the Animal Welfare Act (AniWA) of the Federal food safety and veterinary office, Switzerland. The brains were extracted post-mortem and fixed by immersion in 4% PFA with 15% picric acid (PA) for 24 h. The exact ages of the animals were unknown; they were aged as adults (categorizable in the middle age life stage) by experienced veterinarian with the following criteria: the closure of the femoral and humeral epiphyseal plate, the body weight, the forearm length and sexual maturity. Four young adult and one middle-aged marmoset from the George Washington University were provided by the Texas Biomedical Research Institute (USA): they were collected at the time of necropsy following euthanasia. All brains were immersion-fixed in 10% buffered formalin immediately at necropsy. After a 5-day period of fixation, brains were transferred into a 0.1 M phosphate buffered saline (PBS, pH 7.4) solution containing 0.1% sodium azide and stored at 4° C. All procedures were approved by the Institutional Animal Care and Use Committee at The George Washington University, and follow the ethical guidelines outlined by the American Society of Primatologists. None of the brains included in this study showed gross abnormalities or pathology on veterinary inspection.

Four prepuberal and four young adult rabbit (*O. cuniculus*) brains come from a stock at the NICO animal facility used in a previous report (65). Animals were deeply anesthetized (ketamine 100 mg/kg - Ketavet, Bayern, Leverkusen, Germany - and xylazine 33 mg/ kg body weight - Rompun; Bayer, Milan, Italy) and perfused intracardially with a heparinized saline solution followed by 4% paraformaldehyde in 0.1 M sodium phosphate buffer, pH 7.4. Brains were then postfixed for 6 hours. All experiments were in accordance with the European Communities Council Directive of 24 November 1986 (86-609 EU) and the Italian law for the care and use of experimental animals (DL.vo 116/92). All procedures carried out in this study were approved by the Italian Ministry of Health (8 october 2009) and the Bioethical Committee of the University of Turin (66).

The brains of four young adult and four middle-aged cats (*F. catus domestica*), four young adult sheep (*O. aries*), four young adult and four middle-aged horses (*E. caballus)* were provided from the University of Padova (tissue samples preserved in the MMMTB are distributed to qualified research centers worldwide, following a specific documented request, according to national and international regulations on protected species - CITES). These brains, obtained post-mortem, were fixed by immersion in 10% buffered formalin and kept in the fixative solution for one month (sheep, cat), and three months (horses). Sheep and horse samples originated from a commercial slaughterhouse and were treated according to the European Community Council directive (86/609/EEC) on animal welfare during the commercial slaughtering process and were constantly monitored under mandatory official veterinary medical care. The definition of each age group was based on the official documentation available, corresponding to each individual ear mark in sheep, and confirmed by teeth examination in horses. Four prepuberal and four middle-aged sheep (*O. aries* - breed: Ile de France) were raised at the Institut National de la Recherche Agronomique (INRA; Nouzilly, Indre et Loire, France; ethical permissions are reported in Ref. 67). Four prepuberal sheep were perfused through both carotid arteries with 2 L of 1% sodium nitrite in phosphate buffer saline, followed by 4 L of ice-cold 4% paraformaldehyde solution in 0.1 M phosphate buffer at pH 7.4. The brains were then dissected out, cut into blocks and post-fixed in the same fixative for 48 h. Four middle-aged sheep were collected 20 min after death and kept in 10% formalin for 1 month.

Four young adult and four aged chimpanzee (*P. troglodytes*) brains were provided by the National Chimpanzee Brain Resource (USA); they were collected post-mortem from Association of Zoos and Aquariums, maintained in accordance with each institution’s animal care guidelines and fixed by immersion in 10% formalin (68). After 10-14 days the brain was transferred in 0.1 M phosphate buffered saline (PBS) with 0.1% sodium azide solution and stored at 4°C. Experiments were conducted following the international guiding principles for biomedical research involving animals developed by the Council for International Organizations of Medical Sciences (CIOMS) and were also in compliance with the laws, regulations, and policies of the “Animal welfare assurance for humane care and use of laboratory animals,” permit number A5761-01 approved by the Office of Laboratory Animal Welfare (OLAW) of the National Institutes of Health, USA.

#### 4.2.2 Tissue processing for histology and immunohistochemistry

After fixation, the whole hemispheres of marmoset, cat, sheep, chimpanzee, and horse brains were cut into coronal slices (1-2 cm thick). The slices were washed in a phosphate buffer (PB) 0.1 M solution, pH 7.4, for 24-72 hours (based on brain size) and then cryoprotected in sucrose solutions of gradually increasing concentration up to 30% in PB 0.1 M. Slices of larger brains (chimpanzee and horse) were further reduced in size to be processed in the cryostat or microtome, by cutting them into two blocks, dorsal and ventral. Then, slices, blocks or the entire hemispheres (mouse, naked mole rat, rabbit) were frozen by immersion in liquid nitrogen-chilled isopentane at −80°C. Before sectioning, they were kept at −20°C for at least 5 hours (time depending on brain size) and then cut into 40 μm thick coronal sections using a cryostat (total numbers of sections/hemisphere are reported in Table S4). Free-floating sections were then collected and stored in cryoprotectant solution at −20 °C until staining.

Sections were used both for histological staining procedures (aimed at defining the overall neuroanatomy and boundaries of the entire hemisphere or amygdala for volume estimations), and for immunocytochemical detection of specific markers (Table S2). Histological analyses were performed on Toluidine blue stained sections (cresyl violet for chimpanzees). For immunohistochemistry, two different protocols of indirect staining were used: peroxidase or immunofluorescence techniques. For 3,3’-diaminobenzidine (DAB) immunohistochemistry, free-floating sections were rinsed in PBS 0.01 M, pH 7.4. Antigen retrieval was performed using citric acid, pH 6.0, at 90°C for 5-45 minutes. After further washing in PBS 0.01 M, the sections were immersed in appropriate blocking solution (1-3% Bovine Serum Albumin, 2% Normal Horse Serum, 0,2-2%Triton X-100 in 0.01M PBS) for 90 minutes at RT. Following, sections were incubated with primary antibodies for 48 hours at 4°C (Table S2). After washing in PBS 0.01 M, sections were incubated for 2 hours at RT with biotinylated secondary antibodies (anti-goat, made in horse, 1:250; anti-mouse made in horse, 1:250; anti-rabbit made in horse, 1:250, Vector Laboratories). Then, sections were washed with PBS 0.01 M, and incubated in avidin–biotin–peroxidase complex (Vectastain ABC Elite kit; Vector Laboratories, Burlingame, CA 94010) for 1 hour at RT. The reaction was detected with DAB, as chromogen, in TRIS-HCl 50 mM, containing 0,025% hydrogen peroxide for few minutes and then washed in PBS 0.01 M. Sections were counterstained with Toluidine blue staining, mounted with NeoMount Mountant (Sigma-Aldrich, 1090160500) and coverslipped.

For immunofluorescence staining, free-floating sections were rinsed in PBS 0.01 M. Antigen retrieval was performed using citric acid at 90°C for 5-30 minutes. After further washes in PBS 0.01 M, sections were immersed in appropriate blocking solution (1-3% Bovine Serum Albumin, 2% Normal Donkey Serum, 1-2% Triton X-100 in 0.01M PBS) for 90 minutes at RT. Then the sections were incubated for 48 hours at 4°C with primary antibodies (Table S2), and subsequently with appropriate solutions of secondary antibodies: Alexa 488-conjugated anti-mouse (1:400; Jackson ImmunoResearch, West Grove, PA), Alexa 488-conjugated anti-rabbit (1:400; Jackson ImmunoResearch, West Grove, PA), cyanine 3 (Cy3)-conjugated anti-goat (1:400; Jackson ImmunoResearch, West Grove, PA), cyanine 3 (Cy3)-conjugated anti-guinea pig (1:400; Jackson ImmunoResearch, West Grove, PA-706-165-148), Alexa 647-conjugated anti-mouse (1:400; Jackson ImmunoResearch, West Grove, PA), Alexa 647-conjugated anti-rabbit (1:400; Jackson ImmunoResearch,West Grove, PA 711-605-152) antibodies for 4 hours at RT. Immunostained sections were counterstained with 4’,6-diamidino-2-phenylindole (DAPI, 1:1000, KPL, Gaithersburg, Maryland USA) and mounted with MOWIOL 4-88 (Calbiochem, Lajolla,CA).

#### 4.2.3. Comparable neuroanatomy of the mammalian species considered

The mammalian brains analyzed here differ in terms of brain size, gyrencephaly, and overall neuroanatomical organization. In addition to interspecies differences, some other variables were present, mostly due to tissue processing: brains of the same species can differ in single individuals and can undergo variable degrees of shrinkage (e.g., temperature during cutting might affect section thickness). Thus, each specimen in the age groups of each animal species can show slightly different numbers of coronal sections covering the entire hemisphere (and consequently the amygdala). To account for this variable, a fixed number of serial coronal sections covering both the whole hemisphere and amygdala was considered in each species, as follows: the total number of sections obtained for each specimen/species were compared, and the specimen with the lower number of sections was used as a reference to reach the same number of sections in all specimens. To do this, sections at the very beginning and at the very end of the brain (in the frontal and occipital lobes) were excluded. The same was done for the amygdala, by excluding sections at the very beginning and very end of the structure. In this way, sections were comparable among all specimens of a single species (the numbers of sections used is reported in Table S4).

#### 4.2.4. Volume measurement and cell counting

*Volume measurement.* To estimate the volume of the brain hemisphere and amygdala region, serial coronal sections at a 480 μm interval apart (1 section out of 12 serial sections; 1 out of 48 in the very large brains of horse and chimpanzee, only for whole hemisphere analysis; see Table S4 for number of sections employed for the analysis) were collected covering the whole hemisphere and amygdala anterior-posterior extension in each species (3 specimen/species at each life stage). The selected sections were stained with Toluidine blue to highlight the overall neuroanatomy (Figs. 1C,D and 4C); the stained sections were scanned using Axio Scan (Zeiss; Oberkochen, Germany) and the whole hemisphere coronal sections and amygdala areas for each specimen were measured using the “Contour Line” tool of ZEN Blue Software (Zeiss; Oberkochen, Germany). The volume was calculated in each specimen by using the following formula:

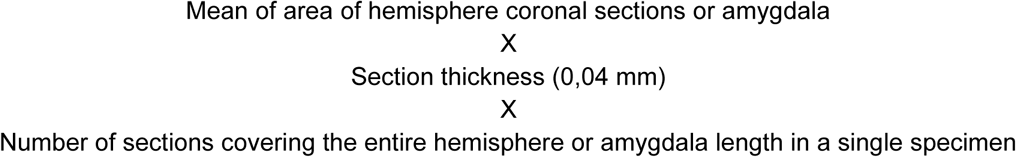

*Cell counting.* To perform the quantitative analysis of both DCX^+^ cells and Ki67^+^ nuclei, serial sections at a 480 μm interval from each other were collected covering the whole amygdala in each species (4 specimen/species at each life stage; Figs. 1 and 4). The counting was done using Neurolucida software (MicroBrightfield, Colchester, VT) as follows: a trained operator performed the tracing of the amygdala area in each section using the “Contour” tool of Neurolucida and all positive stained cells herein (objective: 20x magnification) were counted with markers (Fig. 4A,B) to obtain the DCX^+^ or Ki67^+^ cell density/mm^2^. Cells cut on the superior surface of the section were not considered, to avoid overcounting. The cell soma size was obtained by evaluating the cell soma width (diameter orthogonal to main axis), measured in about 100 cells for each animal species using the Neurolucida ‘measure line’ tool.

The search for possible Ki67^+^/DCX^+^ double-stained cells was performed in 40x magnification confocal fields. For all specimens (aside from chimpanzees), 3 sections from the anterior, middle and posterior part of the amygdala were selected and, in each of them, three microscopic fields in the dorsal, medial and ventral part of amygdala were analyzed.

The percentage of NeuN^+^/DCX^+^ double-stained cells was measured in marmoset, rabbit and cat (3 specimens/species). For each specimen, 3 sections from the anterior, middle and posterior part of the amygdala were selected and, in each of them, three microscopic fields including both type 1, type 2a and type 2b cells were acquired. A total of 27 confocal fields/species and at least 300 cells/species were counted for the analysis (tabulated data can be found in data S1).

The number of sections cut in each hemisphere and those considered for counting procedures in each species are listed in Table S4. A total of 4365 cryostat sections coming from 80 brains were analyzed.

#### 4.2.5. Image acquisition and processing

Images were collected using a Nikon Eclipse 90i microscope (Nikon, Melville, NY) connected to a color CCD Camera, a Leica TCS SP5, Leica Microsystems, Wetzlar, Germany and a Nikon Eclipse 90i confocal microscope (Nikon, Melville, NY). For volume analysis, Zeiss Axio Scan.Z1 and Zen Blue software were used (Zeiss; Oberkochen, Germany). For quantitative analysis of DCX^+^ and Ki67^+^cells, Neurolucida software (MBF Bioscience, Colchester, VT) was used. For both Ki67^+^/DCX^+^, and NeuN^+^/DCX^+^ double-stained cell analyses, “Cell Counter” Plugin of ImageJ software (version 1.50b; Wayne Rasband, Research Services Branch, National Institute of Mental Health, Bethesda, Maryland, USA) was used.

All images were processed using Adobe Photoshop CS4 (Adobe Systems, San Jose, CA) and ImageJ version 1.50b (Wayne Rasband, Research Services Branch, National Institute of Mental Health, Bethesda, Maryland, USA). Only general adjustments to color, contrast, and brightness were made. The percentages of areas occupied by the immature cells (area fraction; the percentage of pixels in the selection that have been highlighted in red; see Fig. 4D) in each of the amygdala subnuclei (and in each of its anterior, middle, and posterior part) were measured by using the Image/Adjust/Threshold function of ImageJ imported from the Neurolucida fields employed for DCX^+^ cell counting.

#### 4.2.6. Statistical and phylogenetic analysis

All graphs and relative statistical analysis were performed using GraphPad Prism Software (San Diego California, USA) using different nonparametric tests: Mann-Whitney test, Kruskal-Wallis test with Dunn’s multiple comparison post-test. p < 0.05 was considered statistically significant. Medians were used as data central measure as previously done for the analysis of INs in the cerebral cortex (15).

Species mean DCX^+^ and Ki67^+^ cell densities were used to perform ancestral character state reconstructions of trait evolution mapped onto the phylogeny. A phylogenetic tree of the species in the sample was downloaded from the TimeTree database (69). The ancestral character state reconstruction was implemented in Mesquite software (version 3.81), using a parsimony model.

To determine the scaling relationships in our dataset we employed least squares regression. All data were log transformed to fit power functions to linear regression, as is standard procedure in comparative studies of neuroanatomy.

## Supporting information

Supplementary Tabulated data

## Acknowledgments

We thank Irmgard Amrein for her generous gift of adult marmoset brains, Ugo Ala for precious advice in statistical analyses, Alessandro Zanone, Eleonora Pintauro, Elaine Miller, and Dustin Howard for technical help in some experimental procedures, and Enrica Boda for stimulating discussion on glial cells.

## Author contributions

Conceptualization: L.B. Methodology: M.G., C.C.S., L.B. Formal Analysis: M.G., N.T., C.C.S. Investigation: M.G., C.L., J.M.G., C.G.F. Resources: C.L., J.M.G., C.G.F., C.C.S., L.B. Data Curation: M.G. Writing - Original Draft: L.B., M.G., C.C.S. Writing - Review & Editing: C.C.S., M.G., J.M.G., L.B. Visualization: L.B., M.G., C.C.S. Supervision: L.B. Funding Acquisition: L.B., C.C.S.

## Data and materials availability

All data needed to evaluate the conclusions in the paper are present in the paper and/or the Supplementary Materials.

