## Supplementary Tables and Figure for "Phylogenetic variation of immature neurons in mammalian amygdala: high prevalence in primate expanded nuclei projecting to neocortex"

### This PDF file includes:

Fig. S1
Tables S1 to S6

### Other Supplementary Materials for this manuscript include the following:

Data S1 (Excel file, submitted as a separate file)

**Figure S1. Comparative neuroanatomy of amygdala extension and subnuclei segmentation in all mammals considered.** For subnuclei segmentation, the entire amygdala anterior-posterior length of each species (from 1,4 to 9,6 mm) was analyzed at 480 µm pace (one serial coronal section out of 12 stained with toluidine blue; cresyl violet in chimpanzees) and matched with: (7, 70, mouse; 71, naked mole rat; 72, 73, rabbit; 28, 74, cat; 75, 76, sheep; 77, 78, marmoset; 46, chimpanzee; 79, horse). The representation of subnuclei (on the right) is shown in three parts of the amygdala (anterior, middle, posterior; see drawings of the correspondent sections on the left), some nuclei having different length. The paralaminar nucleus has been represented only in species wherein it has been previously described (mouse and marmoset). Nuclei of the basolateral complex (BLc, including the paralaminar nucleus) are colored in grey. This interspecies mini-atlas was used to establish the topographical distribution of the $DCX^+$ and $Ki67^+$ cells in both longitudinal (anterior-to-posterior extension, showed in Fig. 6) and coronal axis (lateral-medial and dorsal-ventral position, Fig. 7). Animal species are arranged from top to bottom according to increasing brain size.

Figure S1

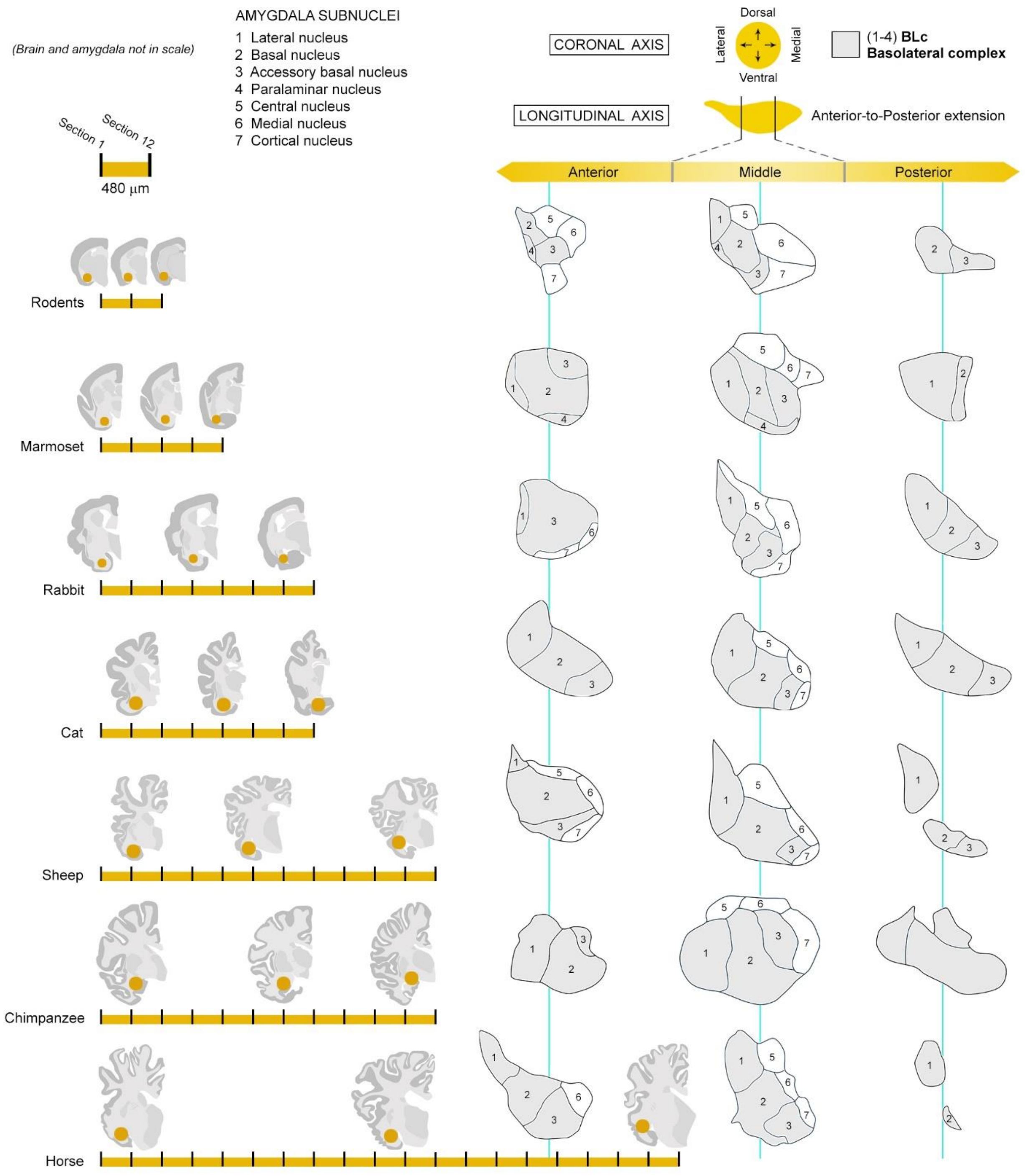
(Brain and amygdala not in scale)
AMYGDALA SUBNUCLEI
1 Lateral nucleus
2 Basal nucleus
3 Accessory basal nucleus
4 Paralaminar nucleus
5 Central nucleus
6 Medial nucleus
7 Cortical nucleus
CORONAL AXIS
Dorsal
Lateral
Medial
Ventral
(1-4) BLc
Basolateral complex
LONGITUDINAL AXIS
Anterior-to-Posterior extension
Anterior
Middle
Posterior
Section 1
Section 12
480 µm
Rodents
Marmoset
Rabbit
Cat
Sheep
Chimpanzee
Horse

**Table S1.** Animals and brain tissues used in this study (4 specimens/age)

| Species | Source | Ages | | Fixation | Fixative | PMI |
|---|---|---|---|---|---|---|
| *Mouse* | (d) | (PP) 10 d | | Immersion | 4% PFA | A few minutes |
| | | (**YA**) 3 m | | Perfusion (IC) | | None |
| | | (MA) 9 m | | | | |
| | | (AG) 15 m | | | | |
| *Naked mole rat* | (f) | (PP) 2 m | | Immersion | | A few minutes |
| | | (**YA**) 2 y | | Perfusion (IC) | | None |
| | | (MA) 10 y | | | | |
| *Marmoset* | (h) | (**YA**) 2.5 y * | | Immersion | 10% formalin | None |
| | (a) | (MA) 5-8 y | 5-8 y<br>5-8 y<br>5-8 y<br>6 y | | 4% PFA+15% PA | 1 hour |
| *Rabbit* | (d) | (PP) 3 m | | Perfusion | 4% PFA | None |
| | | (**YA**) 3 y | | | | |
| *Cat* | (b) | (**YA**) 1.5 y | | Immersion | 10% formalin | 1 hour |
| | | (MA) 6-7 y | 6 y<br>7 y<br>7 y<br>7 y | | | |
| *Sheep* | (e) | (PP) 4 m | | Perfusion (CA) | 4% PFA | None |
| | (b) | (**YA**) 5-6 y | 5 y<br>6 y<br>6 y<br>6 y | Immersion | 10% formalin | 20 minutes |
| | | (MA) 8-10 y | 8 y<br>10 y<br>10 y<br>10 y | | | |
| *Chimpanzee* | (c) | (**YA**) 17-24 y | 17 y<br>18,5 y<br>19,3<br>24 | Immersion | | 14 hours |
| | | (AG) 40-48 y | 40 y<br>41,6 y<br>44,5 y<br>48 y | | | |
| *Horse* | (b) | (**YA**) 3-7 y | 3 y<br>3 y<br>4 y<br>7 y | Immersion | | 1 hour |
| | | (MA) 11-16 y | 11 y<br>15 y<br>15 y<br>16 y | | | |

(a) Institute of Anatomy – University of Zurich; (b) Department of Comparative Biomedicine and Food Science – University of Padova; (c) National Chimpanzee Brain Resource – USA; (d) Neuroscience Institute Cavalieri Ottolenghi (NICO); (e) INRA research center – Nouzilly, France; (f) School of Biological & Chemical Sciences, Queen Mary University of London, London; (h) George Washington University, Washington DC. PMI: postmortem interval; CA, carotid artery; IC, intra-cardiac; PFA, paraformaldehyde solution. Age groups: PP, Prepuberal; PA, Picric acid; YA, Young adult (in bold since a stage present in all species); MA, middle age; AG, aged; *see text; d, days; m, months; y, years. IC, intracardiac; CA, carotid artery. Animal species are arranged from top to bottom according to their increasing brain size. For information regarding ages the Animal Diversity Web (80; available at https://animaldiversity.org/) was used.

**Table S2.** Primary antibodies used in this study

| Antigen | Host | Type | Code | Raised against | Dilution | Source |
|---|---|---|---|---|---|---|
| DCX | goat | polyclonal | SC8066 | Epitope within the last 50 c-terminal amino acids | 1:300-2000 | Santa Cruz Biotechnology |
| | mouse | monoclonal | SC271390 | Amino acids 81-365 mapping at the C-terminus of Doublecortin of human origin | | |
| | rabbit | | AB18723 | Synthetic peptide conjugated to KLH derived from within residues 300 to the C-terminus of Human Doublecortin | 1:1000 | Abcam |
| | | polyclonal | 4604 | Antigenic sequence surrounds amino acid 350 tyrosine of human doublecortin | | Cell Signaling |
| | guinea pig | | AB2253 | Epitope aminoacidic sequence: YLPLSLDDSDSLGDSM | | Merck Millipore |
| Ki-67 | rabbit | | AB15580 | Synthetic peptide | 1:500 | Abcam |
| | mouse | monoclonal | 550609 | Human Ki-67 | | BD Pharmingen |
| PSA-NCAM | mouse | | MAB5324 | Viable Meningococcus group B (strain 355) | 1:700 | Merck Millipore |
| NeuN | mouse | | MAB377 | Purified cell nuclei from mouse brain | 1:300 | |
| Tbr1 | rabbit | polyclonal | AB10554 | KLH-conjugated linear peptide corresponding to 18 amino acids from the N-terminal region of mouse T-box brain protein 1 (Tbr1) | 1:1000 | |
| SOX10 | | | HPA068898 | Immunogen sequence: PHYTDQPSTSQIAYTSLS LPHYGSAFPSISRPQFDY SDHQPSGPYYGHSG | 1:500 | |
| Olig2 | | | AB9610 | Recombinant mouse Olig-2 | | |

**Table S3.** Percentages of areas occupied by $DCX^+$ immature neurons in the amygdala subnuclei

| **Species** | La | Ba (+PL)* | Ab | **BLc** | | Ce | Me | **Ce-Me** | | Co |
|---|---|---|---|---|---|---|---|---|---|---|
| *NMR* | 0 | 0,002 | 0,003 | **0,005** | | 0 | 0 | **0** | | 0 |
| *Mouse* | 0 | 0,004 | 0 | **0,004** | | 0 | 0 | **0** | | 0 |
| *Sheep* | 0,007 | 1,472 | 1,169 | **2,648** | | 0,015 | 0,015 | **0,030** | | 0,110 |
| *Cat* | 0,270 | 0,457 | 0 | **0,727** | | 0,060 | 0,010 | **0,070** | | 0,600 |
| *Rabbit* | 0,097 | 0,052 | 0,205 | **0,285** | | 0,010 | 0,040 | **0,025** | | 0,210 |
| *Horse* | 0,280 | 0,673 | 1,630 | **2,040** | | 0,280 | 0,790 | **0,930** | | 4,820 |
| *Marmoset* | 0,053 | 3,230 | 0,985 | **3,940** | | 0 | 0 | **0** | | 0 |
| *Chimpanzee* | 1,873 | 6,147 | 0,080 | **8,100** | | 0,040 | 0,160 | **0,200** | | 2,700 |

La, lateral nucleus; Ba, basal nucleus (*the PL has been included both for species in which it has been previously described and in the others, wherein it is considered as part of the Ba); Ab, accessory basal nucleus; PL, paralaminar nucleus; BLc, basolateral complex (including the PL); Ce, central nucleus; Me, medial nucleus; Ce-Me, centro-medial nucleus; Co, cortical nucleus. Background colors (dark grey, white, and light grey) identify the three amygdala subdivisions showed in Fig. 7B.

**Table S4.** Number of cryostat sections cut in each hemisphere and used for quantitative analyses in the different animal species and ages

<table>
<tr><th>Species</th><th>Age</th><th>N. of sections in the whole hemisphere</th><th>Whole hemisphere volume analysis</th><th>N. of sections in the whole amygdala</th><th>Cell counting and volume analysis in the amygdala</th></tr>
<tr><td rowspan="4">Mouse</td><td>PP</td><td>96</td><td>8</td><td rowspan="7">36</td><td rowspan="7">3</td></tr>
<tr><td>YA</td><td>144</td><td>12</td></tr>
<tr><td>MA</td><td>180</td><td>15</td></tr>
<tr><td>AG</td><td>144</td><td>12</td></tr>
<tr><td rowspan="3">NMR</td><td>PP</td><td>96</td><td>8</td></tr>
<tr><td>YA</td><td>120</td><td>10</td></tr>
<tr><td>MA</td><td>120</td><td>10</td></tr>
<tr><td rowspan="2">Marmoset</td><td>YA</td><td>696</td><td>58</td><td rowspan="2">60</td><td rowspan="2">5</td></tr>
<tr><td>MA</td><td>492</td><td>41</td></tr>
<tr><td rowspan="2">Rabbit</td><td>PP</td><td>420</td><td>35</td><td>84</td><td>7</td></tr>
<tr><td>YA</td><td>384</td><td>32</td><td>96</td><td>8</td></tr>
<tr><td rowspan="2">Cat</td><td>YA</td><td>720</td><td>60</td><td>96</td><td>8</td></tr>
<tr><td>MA</td><td>516</td><td>43</td><td>84</td><td>7</td></tr>
<tr><td rowspan="3">Sheep</td><td>PP</td><td>864</td><td>72</td><td>132</td><td>11</td></tr>
<tr><td>YA</td><td>1116</td><td>93</td><td>144</td><td>12</td></tr>
<tr><td>MA</td><td>1152</td><td>96</td><td>180</td><td>15</td></tr>
<tr><td rowspan="2">Chimpanzee</td><td>YA</td><td>2304</td><td>48</td><td>144</td><td rowspan="2">12</td></tr>
<tr><td>AG</td><td>2304</td><td>48</td><td>144</td></tr>
<tr><td rowspan="2">Horse</td><td>YA</td><td>2640</td><td>55</td><td>240</td><td>20</td></tr>
<tr><td>MA</td><td>2640</td><td>55</td><td>216</td><td>18</td></tr>
</table>

**Table S5.** Estimation of the total number of $DCX^+$ cells in the amygdala of different mammals (one hemisphere).

| **Species, age** | **N. of serial sections cut in the entire amygdala** | **N. of sections considered** | **Average N. of $DCX^+$ cells in a coronal section of the amygdala** | **Estimation of total $DCX^+$ cells in amygdala** |
|---|---|---|---|---|
| Mouse PP | 36 | 3 | 14 | **504** |
| Mouse YA | | | 2 | **72** |
| Mouse MA | | | 0,25 | **9** |
| Mouse AG | | | 0,08 | **2,8** |
| NMR PP | | | 1 | **36** |
| NMR YA | | | 1 | **36** |
| NMR MA | | | 0 | **0** |
| Marmoset YA | 60 | 5 | 561 | **33.660** |
| Marmoset MA | | | 305 | **18.300** |
| Rabbit PP | 84 | **7** | 113 | **9.492** |
| Rabbit YA | 96 | 8 | 95 | **9.120** |
| Cat YA | 96 | 8 | 285 | **27.360** |
| Cat MA | 84 | 7 | 439 | **36.876** |
| Sheep PP | 132 | 11 | 999 | **131.868** |
| Sheep YA | 144 | 12 | 623 | **89.712** |
| Sheep MA | 180 | 15 | 518 | **93.240** |
| Chimpanzee YA | 144 | 12 | 4858 | **699.552** |
| Chimpanzee AG | 144 | | 3292 | **474.048** |
| Horse YA | 240 | 20 | 1396 | **335.040** |
| Horse MA | 216 | 18 | 409 | **88.344** |

**Table S6.** Estimation of the total number of Ki67$^+$ cells in the amygdala of different mammals (one hemisphere).

| Species, age | N. of serial sections cut in the entire amygdala | N. of sections considered | Average N. of Ki67$^+$ cells in a coronal section of the amygdala | Estimation of total Ki67$^+$ cells in amygdala |
|---|---|---|---|---|
| Mouse PP | 36 | 3 | 261 | **9.396** |
| Mouse YA | | | 9 | **324** |
| Mouse MA | | | 3 | **108** |
| Mouse AG | | | 2 | **72** |
| NMR PP | | | 16 | **576** |
| NMR YA | | | 5 | **180** |
| NMR MA | | | 1 | **36** |
| Marmoset YA | 60 | 5 | 15 | **900** |
| Marmoset MA | | | 8 | **480** |
| Rabbit PP | 84 | 7 | 95 | **7.980** |
| Rabbit YA | 96 | 8 | 52 | **4.992** |
| Cat YA | 96 | 8 | 15 | **1.440** |
| Cat MA | 84 | 7 | 4 | **336** |
| Sheep PP | 132 | 11 | 443 | **58.476** |
| Sheep YA | 144 | 12 | 40 | **5.760** |
| Sheep MA | 180 | 15 | 19 | **3.420** |
| Chimpanzee YA | 144 | 12 | 27 | **3.888** |
| Chimpanzee AG | 144 | | 4 | **576** |
| Horse YA | 240 | 18 | 12 | **2.880** |
| Horse MA | 216 | 20 | 5 | **1.080** |

**Data S1. (separate file)**

All tabulated data are available in a separate Excel file
